# Traits enabling persistence of living Promethearchaeota in marine sediments frozen for over 100 kyr

**DOI:** 10.1101/2025.03.11.642519

**Authors:** Renxing Liang, Tatiana Vishnivetskaya, Elizaveta M. Rivkina, Karen G. Lloyd

**Affiliations:** State Key Laboratory of Geomicrobiology and Environmental Changes, China University of Geosciences (Wuhan), Wuhan, 430078, China; Princeton University, Princeton, NJ, USA; University of Tennessee, Knoxville, TN, USA; Institute of Physicochemical and Biological Problems in Soil Science, Russian Academy of Sciences, Pushchino, Moscow Region, Russia; University of Southern California, Los Angeles, USA 90031

**Keywords:** Promethearchaeota, ancient permafrost, protein repair enzymes, long-term survivability, adaptation

## Abstract

The phylum Promethearchaeota (formerly “Asgard” archaea), the microbial progenitors of all Eukaryotes, are abundant throughout Earth’s subsurface, who have been hypothesized to persist over geological timescales in stable environments with little cell division. We therefore examined the genetic adaptations of Promethearchaeota after being frozen for >100 kyr by comparing metagenome-assembled genomes (MAGs) from the Kolyma Lowland, Siberia, to MAGs from other marine and terrestrial sediments worldwide. We reconstructed 22 Promethearchaeota MAGs from 5 classes (*Heimdallarchaeia*, *Gerdarchaeia*, *Lokiarchaeia, Helarachaeia* and *Thorarchaeia*). Six MAGs from the intracellular DNA fraction were > 70% complete before and after DNA repair, and therefore likely represent living Promethearchaeota that have maintained high DNA integrity under cryogenic conditions through geological time. These 6 MAGs were also over 7 times more abundant than all other Promethearchaeota MAGs based on read recruitment. These permafrost Promethearchaeota MAGs are closely related to other non-permafrost Promethearchaeota lineages at the family or genus level and share metabolic potential and genes for DNA and protein repair with them. This suggests that the ability to survive for >100 kyr in permafrost is a trait that is widespread within the Promethearchaeota. No genes were more prevalent in our permafrost MAGs compared to Promethearchaeota MAGs from other environments. The lack of detectable genetic change since these groups were frozen is consistent with the predicted state of inactivity. Furthermore, although DNA repair mechanisms were present in the Promethearchaeota/Eukaryote lineage before the eukaryotic split, Promethearchaeota protein repair mechanisms emerged after the split, suggesting that adaptations to long term dormancy, or aeonophily, may set modern Promethearchaeota apart from eukaryotes. Collectively, our study expands the known habitat range of many subgroups of Promethearchaeota to ancient marine permafrost and suggests they may have extraordinary long-term survivability under cryogenic conditions through geological time.

## Introduction

Advances in high-throughput sequencing and bioinformatic analysis have led to the explosive increase of metagenome-assembled genomes (MAGs) from numerous uncultivated bacterial and archaeal lineages in recent years (1). This has enabled the discovery of diverse, largely uncultured microbial cells that have been hypothesized to persist for thousands of years or longer without replicating or dying (2). Demonstrating that cells can live this long in a state of low metabolic activity is difficult, since experiments cannot be conducted over such timescales. Due to their ubiquity in long-lived stable environments, the deeply-branching Promethearchaeota lineages (formerly the Asgard archaea superphylum) (3) are likely to persist over such timescales (4) . Since the first Lokiarchaeota MAG was reported in 2015 (5), many Promethearchaeota lineages (6, 7) have been recovered globally from diverse environments including hydrothermal vent sediments (8–10), deep-sea sediments (11–13), costal marine sediments (14–16), brackish and saline lake sediments (17), hot springs (9), and terrestrial subsurface sediments (9). Recently, Promethearchaeota have been cultivated in laboratories, enabling the study of their traits directly (18–20). However, laboratory experiments alone are incapable of testing whether these organisms survive for tens of thousands of years or longer. To run a many-thousands of years experiment on these organisms requires finding them in a location with no movement of cells or nutrients into or out of it over thousands of years, such as ancient permafrost (21). Recently, several Lokiarchaeota lineages were identified by 16S rRNA amplicon sequencing from deep submarine permafrost that has remained frozen since it was inundated by sea water during the Holocene marine transgression along the Siberian Shelf (22). However, this work did not investigate whether these Lokiarchaeota MAGs were likely to be from living cells, nor was metagenomic sequencing deep enough to assess the diversity of Promethearchaeota genomes present or the traits that may have enabled them to persist in a non-growing state.

The phylogenomic analyses of Promethearchaeota based on concatenated single-copy genes has challenged the three domains of life and shown that the Heimdallarchaeoal lineages within the Promethearchaeota are the closest living relatives of eukaryotes (5–7, 9, 17, 23). However, it is unknown whether Promethearchaeota and eukaryotes share strategies for long-term persistence in a maintenance state that prioritizes repair of biomolecules like DNA and proteins. *RecA*, involved in the repair of double strand DNA breaks (24, 25), and the universal DNA mismatch repair genes (*MutS* and *MutL*) have a shared ancestry between the Promethearchaeota and the eukaryotes (26, 27). Unlike these well-studied mechanisms for DNA repair, the distribution and evolutionary history of ancient and highly conserved protein repair enzymes remain largely unexplored particularly from Promethearchaeota. Since protein repair is likely to taken on outsized importance in tolerance to long-term starvation, over the preservation of other biomolecules (28), protein repair mechanisms may play a prominent role in adaptations to long-term persistence. Protein-L-isoaspartyl (D-aspartyl) *O*-methyltransferase (*PIMT*) is a universal enzyme for the repair of isomerized or racemized aspartyl residues in damaged proteins(29), and the conserved methionine sulfoxide reductases (*MsrA* and *MsrB*) are thioredoxin-linked enzymes that convert methionine sulfoxide to methionine under oxidative stresses (30). These are good candidates as traits for long-term surviving microorganisms through geological time.

Reconstruction of MAGs from ancient permafrost has been challenging due to extremely low biomass and the severe damage of genomic DNA in the dead cells that share the space with the live ones (31, 32). Despite the accrued DNA damage, coupling DNA repair with metagenomics has successfully reconstructed many MAGs from frozen sediments of various geological ages of 26-120 kyr (33). The recovery of a Heimdallarchaeia MAG, from the Promethearchaeota phylum, from one depth within the Kon’kovaya suite (33) prompted us to combine the DNA repair protocol and ultra-deep sequencing to further explore the genomic diversity of Promethearchaeota lineages from various depths of this marine formation that has remained continuously frozen for the past 100-120 kyr. The lack of abiotic racemization of amino acids, as well as the presence of high molecular weight DNA and redox-activated cells in ancient Siberian permafrost suggest that living cells persist there (33). We investigated whether these Promethearchaeota constitute some of this living biomass, as shown by having intact genomes that do not have a higher completeness after DNA repair. We investigated whether the cells of Promethearchaeota that have survived in permafrost for 100 kyr have genetic differences from those present in marine sediments, suggesting that special adaptations are required to survive this geological timescale. Alternatively, they may not have appreciable genetic differences from other Promethearchaeota, which would suggest that the trait of longevity is widespread in this clade. We then performed examined potential adaptive traits for long term survival by exploring their protein repair enzymes (*PIMT, MsrA,* and *MsrB*) to provide new insights into the evolutionary relationship between Promethearchaeota and eukaryotes.

## Material and methods

### Sampling of ancient permafrost sediments

The sampling site is located at Cape Chukochii near the East Siberian Sea coast (70°05’N, 159°55’E). A ∼22 meter long vertical core across a long chronosequence of ancient permafrost (26-120 kyr) (Figure S1) was collected in August, 2017(33). This study focused on the Kon’kovaya suite (Figure S1) that represents the marine horizon situated between non-saline terrigenous sediments(21). This layer of marine permafrost was deposited during a marine transgression and later became frozen around 100-120 kyr ago during the Middle Pleistocene(34). The details of the samples collected and the geochemical characterization has been fully described in our previous study(33).

### DNA extraction and metagenomic sequencing after DNA repair

Sediments from the Kon’kovaya suite (Figure S1) at three different depths (13.4, 14.8 and 18.3 m) were selected for DNA extraction and metagenomic sequencing. Total DNA was extracted from 10 g of sediment using DNeasy PowerMax soil kit (Qiagen, Carlsbad, CA) according to the manufacturer’s procedures. A blank control with sediment-free reagents was incorporated to monitor any potential contamination during the extraction processes. The yield of DNA was quantified by Qubit fluorometer (Invitrogen, CA) and no detectable DNA was observed from the blank control.

Due to the possibility of severe damage in DNA from the ancient marine deposits(33), DNA repair procedures were carried out prior to sequencing according to previously published procedures(33). The PreCR™ Repair Mix (New England Biolabs, MA, USA) contains a variety of enzymes that can repair many types of DNA damage (deaminated cytosine, apurinic/apyrimidinic sites, thymine dimers, nicks and gaps) with the exception of fragmentation and protein-DNA crosslinks. The reaction for DNA repair was performed following the standard protocol provided by the manufacture. Briefly, the PreCR Repair Mix (1 µl) was generally mixed with damaged DNA samples (up to 50 ng) and incubated at 37°C for 20 minutes. The PreCR-repaired DNA was then converted to Illumina sequencing libraries using the Nextera DNA Flex Library Prep kit (Illumina, CA) according to the manufacturer’s instructions. The three metagenomic libraries (13.4, 14.8 and 18.3 m) were sequenced on Illumina NovaSeq 6000 System using the S Prime (SP) Flow Cell. A total of ∼1.5 billion pair-end reads (2×151 nt) were generated from three metagenomes.

### Metagenomic assembly and genome reconstruction

Illumina sequencing adaptors and low quality sequences were removed using fastp v.0.12.6(35) with the following cutoffs (length <50 nt and Phred scores < 30). Given the similarity of microbial community at different depths of the marine horizon, a co-assembling strategy was performed with MEGAHIT v1.1.4(36) using paired-end mode with the settings of *k*-min = 27, *k*-max = 137, *k-* step = 10. The input data for assembly included all the quality-filtered reads from three metagenomes (13.4, 14.8 and 18.3) from this work and another four metagenomes from 14.8 m sample (14.8_iDNA, 14.8_eDNA, 14.8_iDNA_PreCR and 14.8_eDNA_PreCR) from our previous study(33). The co-assembled contigs (> 1.5 kb) were binned using the default settings in the “Binning module” in MetaWRAP v0.8(37) by integrating three different tools, namely MetaBAT v2.12.1(38), MaxBin v2.0(39) and CONCOCT v1.1.0(40). The generated MAGs were refined with the “Bin_refinement module” in MetaWRAP v1.1(37) and further re-assembled with the “Reassemble_bins module” in MetaWRAP v1.1(37). The quality of the reassembled MAGs were assessed with CheckM v1.0.11(41) and all potential Promethearchaeota MAGs were selected for subsequent analyses.

### Taxonomic classification of Promethearchaeota MAGs

The initial taxonomic classification of all Promethearchaeota MAGs was determined using Genome Taxonomy Database Toolkit (GTDB-Tk v 0.3.0)(42) based on 122 archaeal marker genes. A phylogenetic tree was constructed from the concatenated marker genes with FigTree implemented in GTDB-Tk v1.3.0(42). To better identify the Promethearchaeota MAGs (>70% completeness and < 10% contamination), closely related genomes were retrieved from NCBI database (accessed in October, 2020) for further phylogenomic analysis. The sequences of 16 single-copy ribosomal proteins were extracted and aligned with MUSCLE v3.8.31 in Anvi’o v5.2(43). The phylogenetic tree was built with RAxML v8.1.17(44) based on concatenated alignment with the PROTGAMMAILGF and 1000 bootstraps. Additionally, nearly full-length 16S rRNA genes associated Promethearchaeota lineages were assembled from metagenomic reads phyloFlash v3.4(45). Furthermore, 16S rRNA genes found in all Promethearchaeota MAGs were retrieved using Anvi’o 5.2(43). The alignment of all 16S rRNA gene sequences were performed with MUSCLE v3.8.31(46) and the Maximum Likelihood tree was inferred using the Tamura-Nei model using MEGA v7.0.20(47). All finalized phylogenetic trees were edited and visualized using the online iTOL tool(48).

### Metabolic annotation and other analyses

All Promethearchaeota MAGs (>50% completeness and < 10% contamination) were annotated using Prokka v1.13(49) and DFAST tools(50) against TIGRFAM and COG databases. Furthermore, all predicted genes from each MAG were identified by searching against NCBI nr database using “BLASTP” with the default settings (e-value: 1e-5). The taxonomic novelty of Promethearchaeota MAGs was further determined based on average amino acid identity (AAI) calculated using CompareM v0.1.2 (https://github.com/dparks1134/CompareM). The distribution of Promethearchaeota lineages from 13.8, 14.8 and 18.3 m was determined by the relative abundance of each MAG calculated using the “Quant_bin” module in MetaWRAP v1.1(37). Comparative genomic analyses of MAGs from ancient permafrost and other environments were performed following the pangenomics workflow installed in the Anvi’o software(43).

### Assessment of the genomic DNA damage of Promethearchaeota MAGs

Metagenomes from intracellular (iDNA) and extracellular DNA (eDNA) fractions extracted from the 14.8 m sample have been published in our previous study(33). These metagenomes were generated from both iDNA and eDNA fractions with (14.8eDNA_preCR and 14.8iDNA_preCR) and without DNA repair (14.8eDNA and 14.8iDNA)(33). Such unique datasets allowed us to assess the impact of DNA repair on genomic completeness by mapping each Promethearchaeota MAG back to the individual metagenome with and without DNA repair(33). If preserved DNA records past, dead microbial populations, both iDNA and eDNA fractions would have been subjected to damage over geological time. Therefore, DNA repair would enable sequencing of damaged DNA and thereby increase the completeness of MAGs constructed from both iDNA and eDNA fractions(33). By contrast, the DNA repair would show no significant improvement for living microorganism that can constantly repair DNA under frozen conditions over geological time(33, 51). Therefore, the reads from four metagenomes (14.8_iDNA, 14.8_eDNA, 14.8_iDNA_PreCR and 14.8_eDNA_PreCR) were individually mapped to each Promethearchaeota MAG with BWA v0.7.15 implemented in MetaWRAP v0.8 using the “strict” option (no mismatches)(37). The reads mapped to each MAG were assembled using SPAdes v3.13.0(52) in MetaWRAP v0.8(37). The completeness of MAGs was compared in metagenomes derived from both the iDNA and eDNA with and without DNA repair. Since the MAG completeness was only based on the presence of single-copy genes as determined by CheckM v1.0.11(41), the genomic size (actual length in Mb) of each MAG was also compared in order to better show the impact of DNA repair on genomic recovery from each Promethearchaeota lineage.

### Phylogenetic analyses

The *PIMT*, *MsrA* and *MsrB* genes identified from Promethearchaeota MAGs were searched against the NCBI nr database using BLASTP with the default settings. Representative sequences from archaea, bacteria and eukaryotes were retrieved for phylogenetic analyses. All amino acid sequences were aligned using MUSCLE v3.8.31(53) with the default settings. Maximum-likelihood trees were constructed using IQ-TREE v2.0.6(54) with 1000 times ultrafast bootstrapping and the best-fit models determined for *PIMT* (LG+R7), *MsrA* (LG+I+G4) and *MsrB* (LG+R4). The phylogenetic trees for RNA polymerase were also constructed with IQ-TREE v2.0.6(54) using the LG + C60 + G4 + F model for comparison purposes.

To infer the phylogenetic relationship between eukaryotes and Promethearchaeota, 35 single-copy markers(23) from all Promethearchaeota MAGs, and from other selected archaeal, bacterial, and eukaryotic genomes were used for the construction of phylogenetic tree of life. Sequences were aligned with MAFFT-L-INS-I(55) and trimmed using the BLOSUM30 model in BMGE 1.12(56). The maximum-likelihood tree was built with the 35-gene concatenated dataset using IQ-Tree2(54) under the LG + C60 + G4 + F model with 1000 ultrafast bootstraps as suggested previously(23). Furthermore, both maximum likelihood and Baysian analyses were performed based on the concatenation of 33 conserved proteins by excluding two of the largest RNA polymerase subunits. The two RNA polymerase subunits were excluded because they represent the longest universal proteins that might have a strong biased effect on the concatenated alignment of 35 genes to favor the traditional Woese tree of life(57). The Bayesian analyses were performed under the CAT + GTR + G4 model in PhyloBayes-MPI 1.8(58) to account for cross-site compositional heterogeneity(23). Two independent Markov chain Monte Carlo (MCMC) chains were run and a consensus tree with acceptable coverage (maxdiff <0.3) was obtained using the bpcomp program(58). All inferred phylogenetic trees were edited and visualized using the online iTOL tool(48).

### Data availability

All raw sequences from the metagenomes were deposited in Sequence Read Archive under the BioProject PRJNA680161 with accession numbers SAMN18945478, SAMN18945479, SAMN18945480. All genome sequences have been made publicly available at NCBI under the accession numbers (SAMN18946025-SAMN18946039).

## Results

### Genomic recovery of diverse Promethearchaeota from ancient permafrost

By sequencing the post-repair DNA fraction on the Illumina NovaSeq platform, 413.7, 444.7 and 374.9 million quality-filtered reads were generated from the 13.9, 14.8 and 18.3 m samples, respectively. We recovered 22 Promethearchaeota MAGs from the classes Helarchaeia (n=1), Lokiarchaeia (n=16), Thorarchaeia (n=3), Heimdallarchaeia (n=1) and *Gerdarchaeia* (n=1) (Table S1 and S2). A total of 15 Promethearchaeota MAGs with medium- to high-quality (>50% completeness and <10% contamination) were selected for phylogenetic and other analyses (Table S1). Promethearchaeota MAGs were identified at three depths across the Kon’kovaya suite marine horizon, with no clear depth distribution of abundances (Figure S2). The vertical distribution of Promethearchaeota lineages from the three representative depths indicate that most MAGs (11 out of 15) have the highest relative abundance in the deepest layer at 18.3 m (Figure S2).

The taxonomic placements of the Promethearchaeota MAGs were further confirmed based on phylogenetic analyses of concatenated ribosomal proteins (Figure 1), 16S rRNA gene (Figure S3), and 122 archaeal single-copy makers (Figure S4). Gerdarchae*ia*_Ch1_17_bin89 and Heimdallarchae*ia*_Ch1_17_bin51 belong to Gerdarchae*ia* and Heimdallarchae*ia*, respectively (Figure 1). Gerdarchae*ia* _Ch1_17_bin89 is likely the same Promethearchaeota MAG previously reported from the 14.8 m of the same horizon (33) based on the high ANI value (>99.9% ). Gerdarchae*ia* _Ch1_17_bin89 showed high AAI values (42.9-84.9%) and similarity of 16S rRNA genes (up to 93.3%) when compared to the other Gerdarchaeota MAGs (Figure S5). By contrast, the Heimdallarchae*ia*_Ch1_17_51 exhibited much lower AAI values (43.4-64.4%) compared to other known Heimdallarchae*ia* MAGs (Figure S5). Thorarchae*ia*_Ch1_17_bin66 was closely related to Thorarchae*ia*_AB_25 (AAI, 83.7%) (Figure S6). The 12 Lokiarchae*ia* MAGs were highly diverse and scattered into three major subgroups (Figure 1). Two monophyletic groups (Lokiarchae*ia*_Ch1_17_bin106 and Lokiarchae*ia*_Ch1_17_bin24; Lokiarchae*ia*_Ch1_17_bin12 and Lokiarchae*ia*_Ch1_17_bin22) were distantly related to other known Lokiarchae*ia* lineages. The AAI values of the 12 Lokiarchae*ia* MAGs ranged from 46.3% to 87.2% compared to their close relatives (Figure S7).

**Figure 1.**
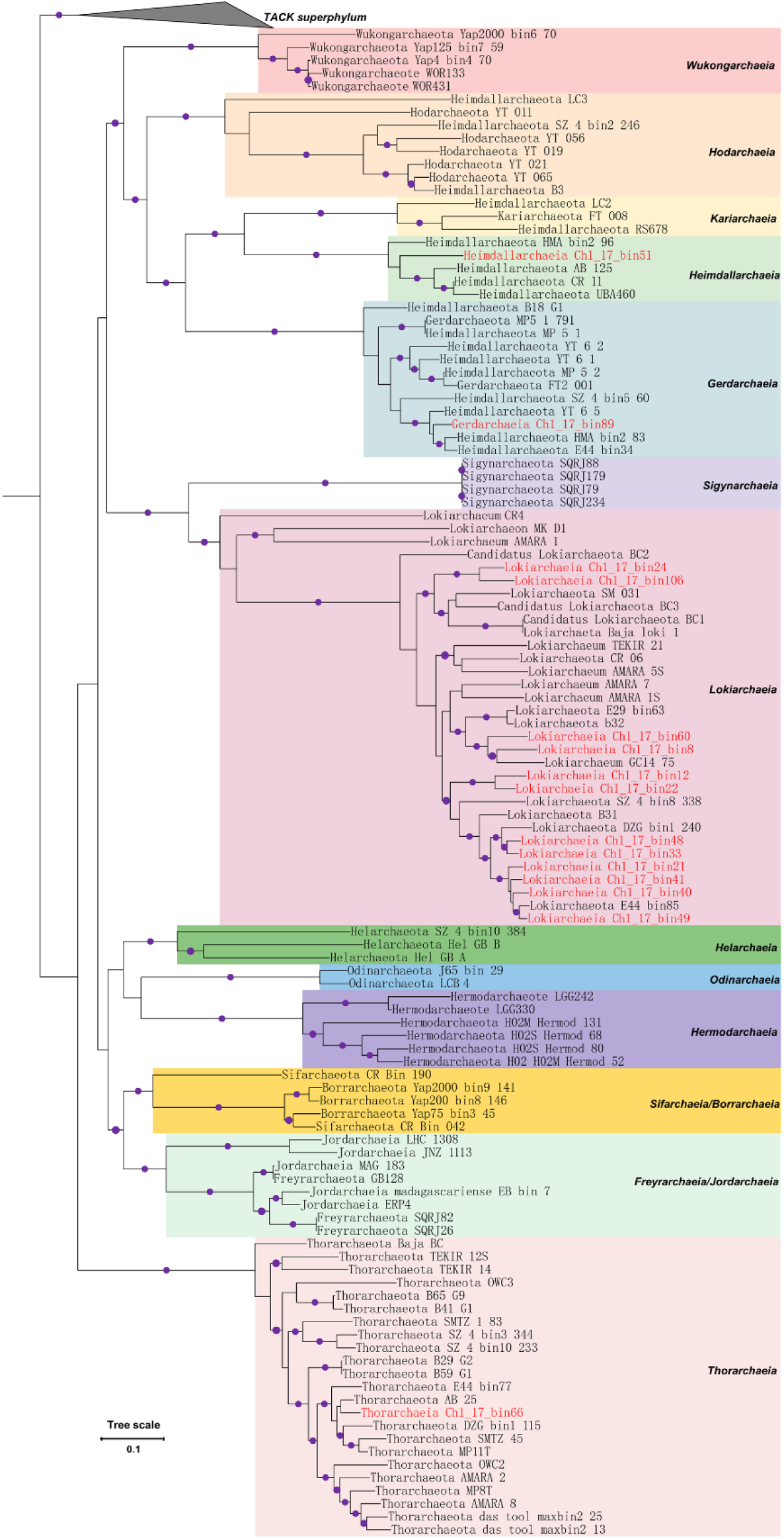
Phylogenetic tree of Promethearchaeota MAGs and their close relatives from diverse environments. The maximum-likelihood phylogenomic tree was constructed based on up to 16 concatenated ribosomal proteins. The purple dots represent bootstrap values > 70% (bootstrap values were generated from 1000 iterations). The TACK group was used as an outgroup. The scale bar corresponds to 0.1 substitutions per amino acid position.

### DNA repair and improved recovery of MAGs

Genomic DNA damage was inferred by comparing the impact of laboratory-based DNA repair on completeness of each Promethearchaeota MAG mapped to individual metagenomes generated from iDNA and eDNA fractions in the 14.8 m sample with and without DNA repair (Figure 2). The genome completeness (up to 26.47%) and size (up to 0.89 Mb) were extremely low for all Promethearchaeota MAGs from the eDNA fraction in the absence DNA repair (Figure 2 and Table S3-S4). However, DNA repair dramatically increased the completeness (2.8-92.05%) and size (0.03-3.4 Mb) of all MAGs from the eDNA fraction (Figure 2 and Table S3-S4). Furthermore, DNA repair also increased the genome size and completeness of most Promethearchaeota MAGs recovered from the iDNA fraction (Figure 2 and Table S3-S4). Notably, six Promethearchaeota MAGs (Gerdarchaeia_Ch1_17_bin89, Lokiarchaeia_Ch1_17_bin48, Lokiarchaeia_Ch1_17_bin12, Lokiarchaeia_Ch1_17_bin22 Lokiarchaeia_Ch1_17_bin41 and Lokiarchaeia_Ch1_17_bin49) exhibited no or slight deviations from the red theoretical line indicating little to no DNA damage (Figure 2).

**Figure 2.**
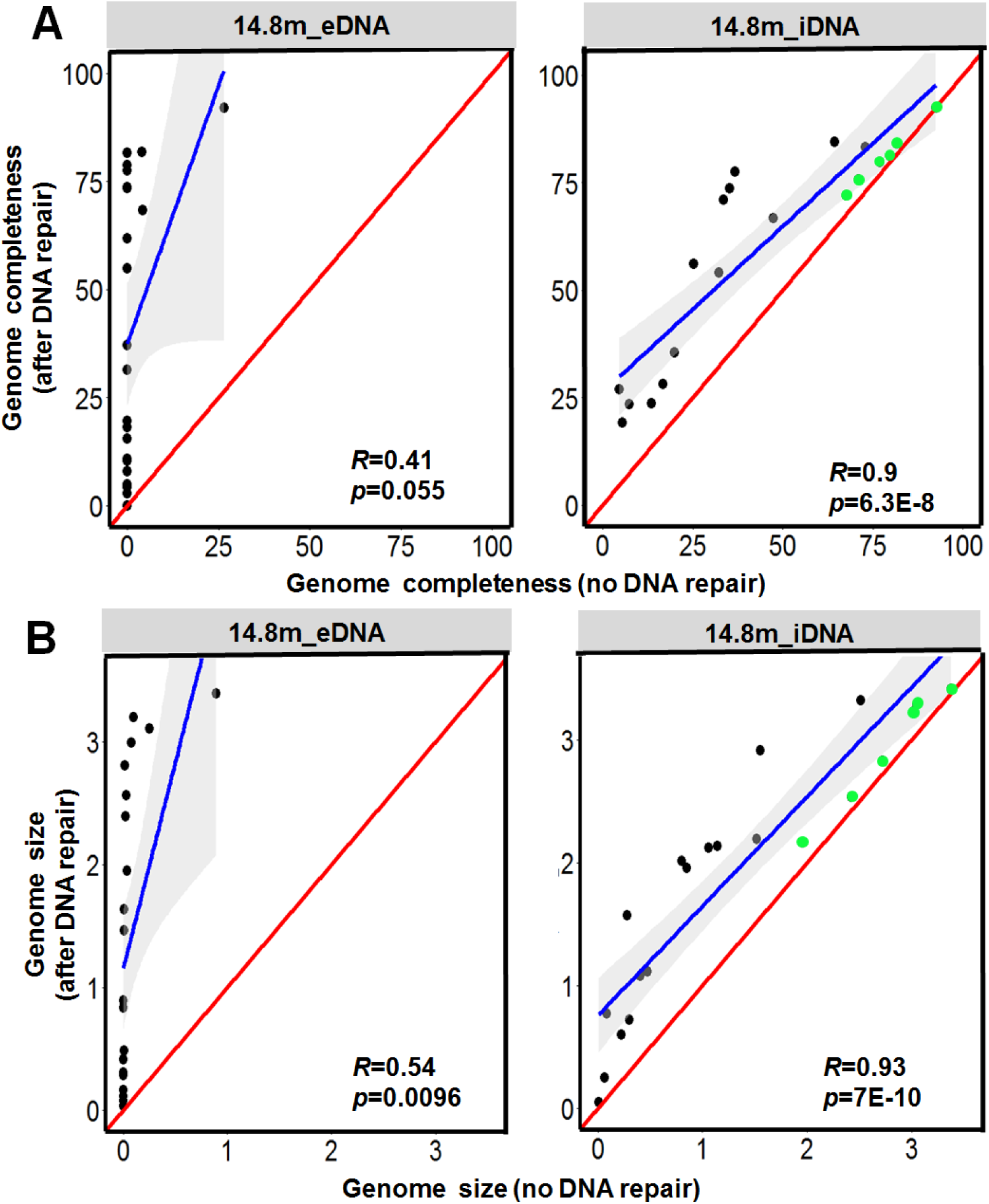
Effect of DNA repair on genome completeness (A) and size (B) of all Promethearchaeota MAGs recovered from individual metagenomes derived from iDNA and eDNA fractions with and without DNA repair. The completeness was determined based on the presence of single-copy genes whereas genomic size refers to the actual length (Mb) the recovered MAGs. The blue line refers to the regression line from the correlation between MAGs completeness with and without DNA repair. The Pearson coefficient (*R*) and p-value (*p*) are shown in each plot. The red line indicates the theoretical line by assuming no DNA damage and thus no positive effect on the MAGs completeness. The green dots represent the MAGs that DNA pair showed minimal impact on the completeness of Promethearchaeota genomes.

### Metabolic reconstruction from Promethearchaeota MAGs

The Promethearchaeota MAGs from permafrost harbor genes involved in the metabolism of various carbohydrates such as polysaccharides (cellulose, starch and xylene) and monosaccharides (glucose, galactose and xylose) (Figure 3). The genes encoding the enzymes of the Embden-Meyerhof-Parnas glycolysis pathway were identified from all Promethearchaeota MAGs with the exception of two Lokiarchaeia MAGs (Lokiarchaeota_Ch1_17_bin60 and Lokiarchaeia_Ch1_17_bin106) (Figure 3), which yielded genes encoding the enzymes of the pentose phosphate glycolysis pathway. Proteolytic enzymes from the families of cysteine peptidases, metallopeptidases, and serine peptidase were detected in most Promethearchaeota MAGs (Figure 3). The genes encoding aldehyde ferredoxin oxidoreductase (AOR), 2-ketoisovalerate ferredoxinoxidoreductase (VOR), and indolepyruvate ferredoxin oxidoreductase (IOR) were present in nearly all MAGs except for Lokiarchaeia_Ch1_17_bin33 and Lokiarchaeia_Ch1_17_bin106. Furthermore, genes involved in the generation of lactate (lactate dehydrogenase), formate (pyruvate formate lyase) and acetate (pyruvate ferredoxin oxidoreductase and ADP-forming acetyl-CoA synthetase) were present in nearly all Promethearchaeota genomes (Figure 2).

**Figure 3.**
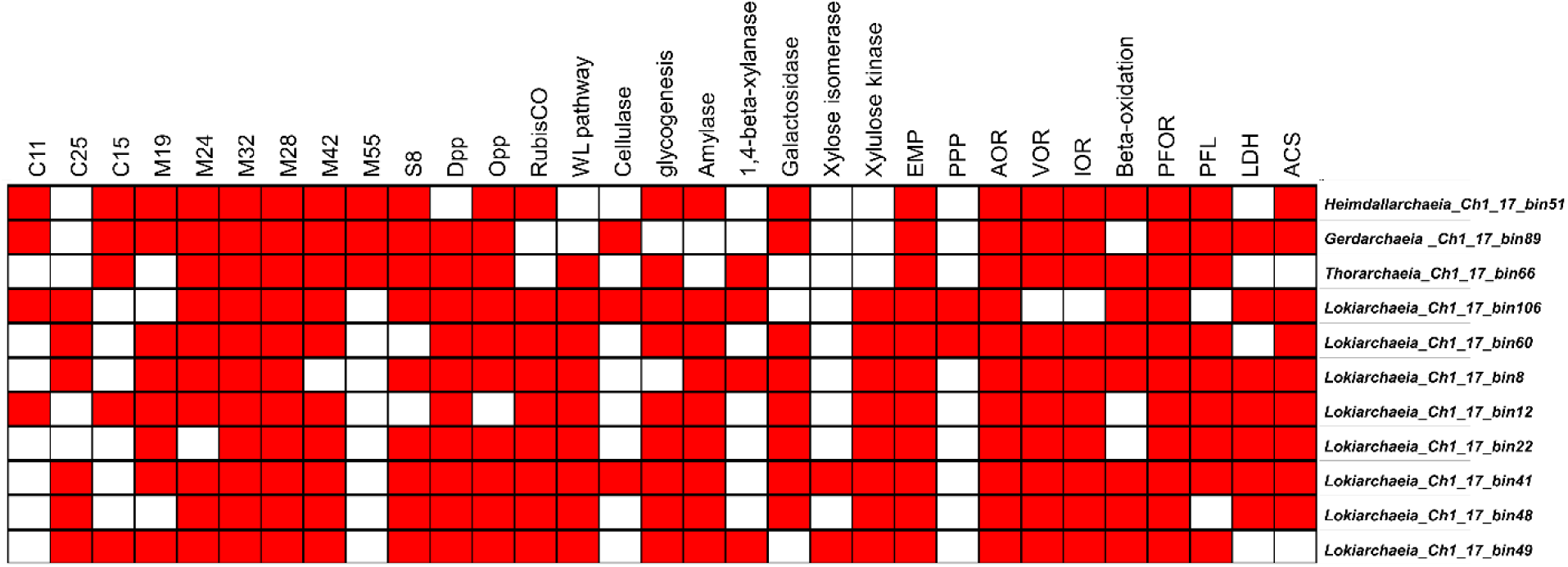
Presence of key functional genes (red) in Promethearchaeota MAGs involved in the metabolism of peptides and carbohydrates. Abbreviations: C11 (Clostripain), C25 (gingipain), C15 (pyroglutamyl-peptidase), M19 (membrane-bound dipeptidase), M24 (Methionyl aminopeptidase), M32 (carboxypeptidase), M28 (aminopeptidase), M42 (metallopeptidase), M55 (DppA aminopeptidase family), S8 (serine peptidase), EMP (Embden-Meyerhof pathway), PPP (pentose phosphate pathway), Aldehyde ferredoxin oxidoreductase (*AOR*), 2-ketoisovalerate ferredoxin oxidoreductase (*VOR*), Indolepyruvate ferredoxin oxidoreductase (*IOR*), *PFOR* (pyruvate:ferredoxin oxidoreductase), PFL (pyruvate formate lyase *L*), LDH (Lactate dehydrogenase), Acetyl-CoA (ADP-forming) synthetase (*ACS*), Wood–Ljungdahl pathway (WL pathway). Note: The RuBisCO pathway refer to the identification of ribulose-1,5-bisphosphate carboxylase and glycogenesis indicates that glycogen synthase and glycogen branching enzyme were identified in the genome.

### Evolutionary history of protein repair enzymes

The PIMT phylogenetic tree (115 bacteria, 21 eukaryotes and 94 archaea) clearly revealed a three-domain topology (Figure 4) that favors the Woese hypothesis. Furthermore, the genes encoding PIMT were identified from all phyla (Heimdallarchaeia, Lokiarchaeia, Thorarchaeia, Helarchaeia and Odinarchaeia) within the Promethearchaeota (Figure 3 and 4). The Promethearchaeota lineages are clustered together with that of the TACK kingdom (Bathyarchaeia and Verstraetearchaeia) as sister groups (Figure 4). In this regard, the PIMT in Promethearchaeota share the same ancestral archaeal lineage with that of the TACK kingdom instead of eukaryotes. By contrast, the eukaryotes form a monophyletic group that is separated from the bacterial and archaeal clades (Figure 4). Unlike the conserved *PIMT* gene, the *MsrA* and *MsrB* genes from different Promethearchaeota phyla were dispersed within different branches of bacterial lineages (Figure 5). Notably, none of the Promethearchaeota genes showed a close phylogenetic relationship to eukaryotes. Furthermore, the structurally distinct enzymes encoded by the *MsrA* and *MsrB* also showed different evolutionary histories between individual Promethearchaeota phyla and bacterial phyla. For instance, the *MsrA* from Lokiarchaeia clustered together with Verrucomicrobia whereas *MsrB* fell within the Spirochaetes and Bacteroidetes (Figure 5).

**Figure 4.**
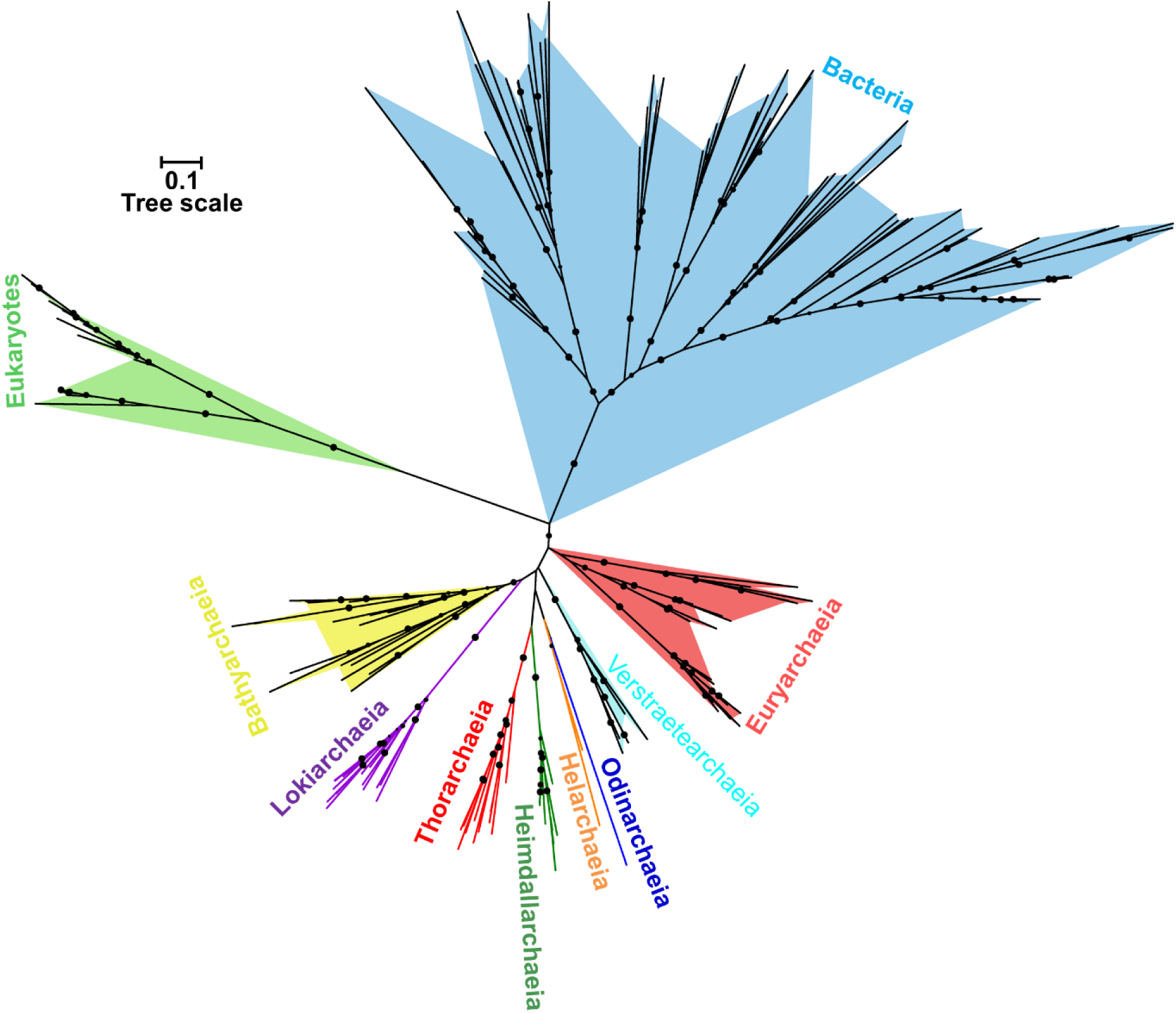
Maximum-likelihood tree of PIMT constructed using IQtree with the LG+R7 model. The black dots represent bootstrap values > 70% (bootstrap values were generated from 1000 iterations). The scale bar corresponds to 0.1 substitutions per amino acid position.

**Figure 5.**
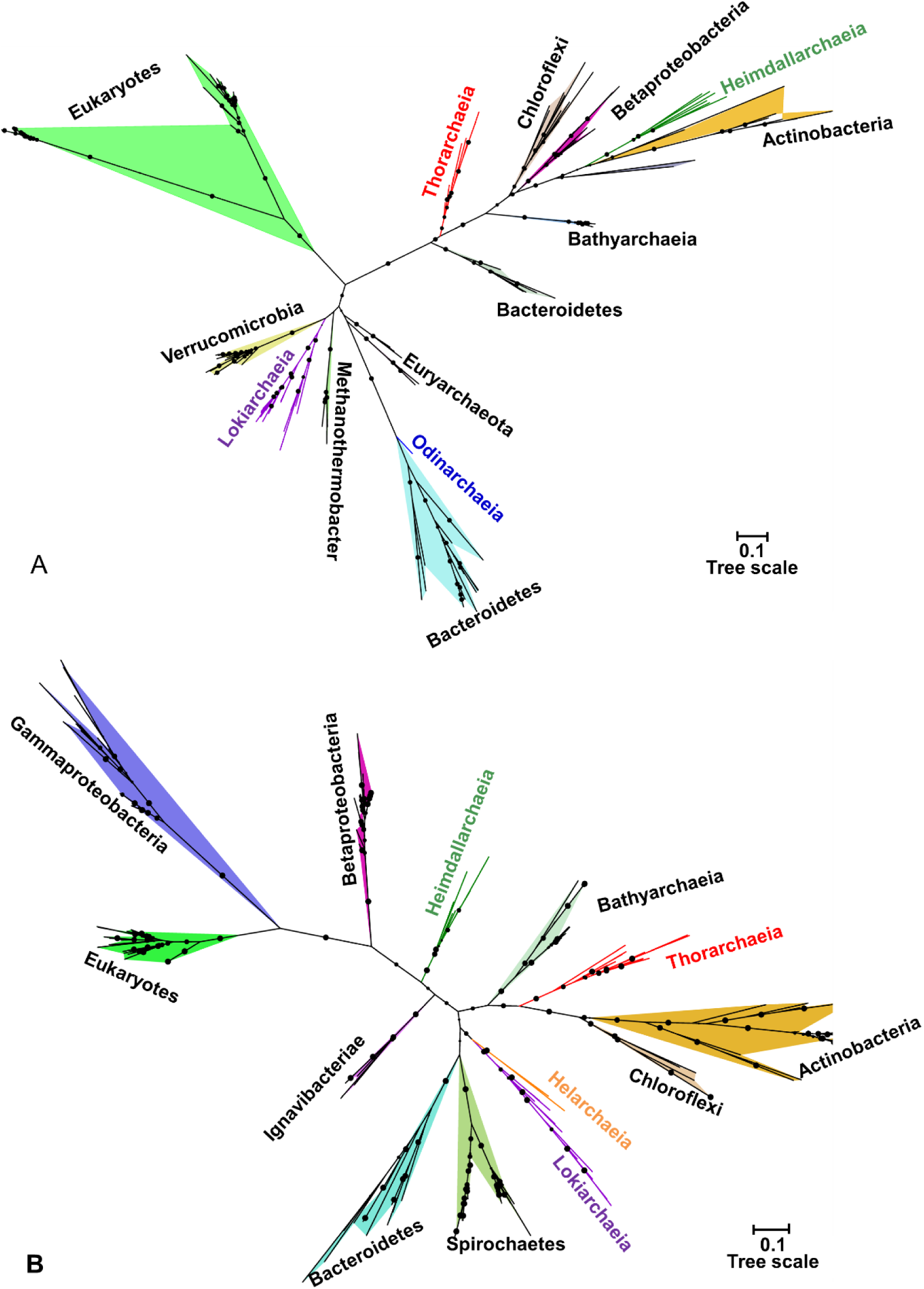
Maximum-likelihood phylogenetic tree for MsrA (A) and MsrB (B) inferred with LG+I+G4 and LG+R4 model, respectively. The black dots represent bootstrap values > 70% (bootstrap values were generated from 1000 iterations). The scale bar corresponds to 0.1 substitutions per amino acid position.

### Evolutionary relationship between Promethearchaeota and eukaryotes

To examine whether the addition of permafrost genomes helps delineate the phylogenetic relationship between Promethearchaeota lineages and eukaryotes, we incorporated an expanded inventory of Promethearchaeota MAGs (87) from permafrost (n=15, this study) and non-permafrost environments to construct a phylogenetic tree from a concatenated alignment of 35 conserved genes (Figure S12). Our results indicated that both maximum-likelihood and Bayesian analyses based on concatenated proteins provided robust support (posterior probability (PP) =1; bootstrap support (BS) = 100) for a close relationship between eukaryotes and the clade containing Heimdallarchaeia, Gerdarchaeia, Kariarchaeia and Hodarchaeia (Figure 6). However, all other Promethearchaeota lineages were more distantly related to eukaryotes (Figure 6).

**Figure 6.**
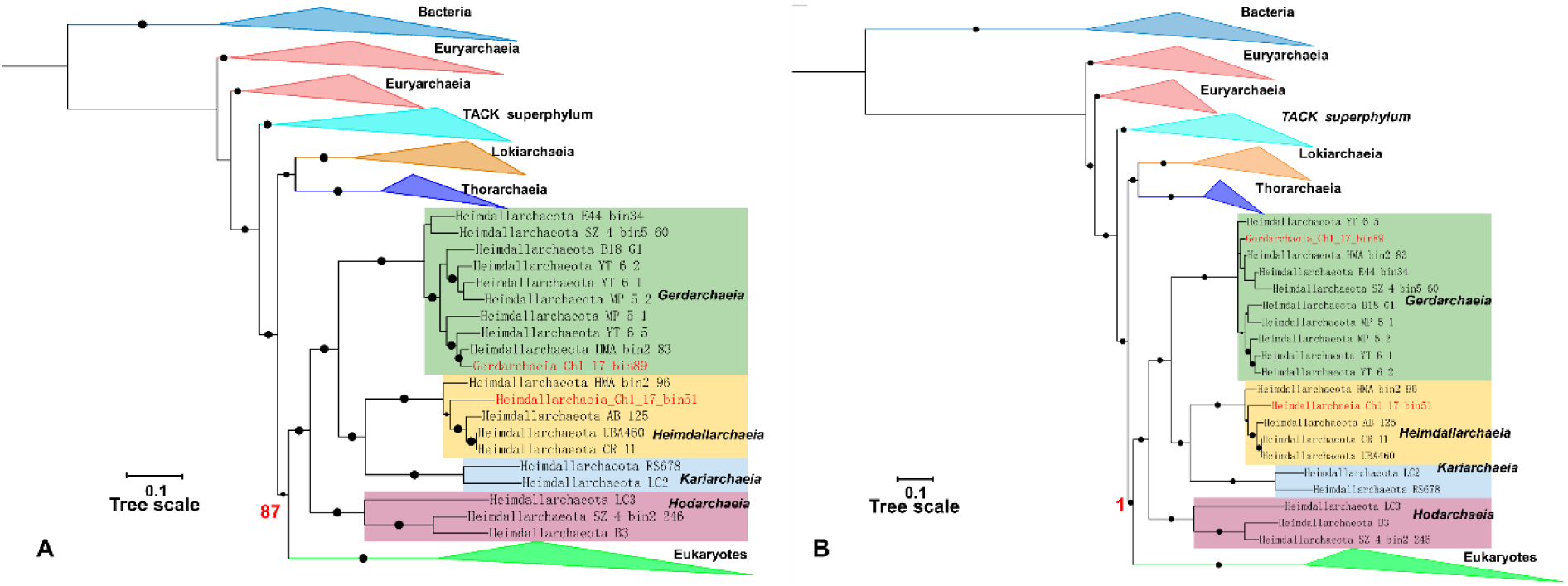
Phylogenetic trees reconstructed using Maximum likelihood analysis under LG +C60 +F+G4 model (A) and Bayesian inference under the CAT + GTR + G4 model (B) based on the concatenation of 33 universal proteins. The black dots on nodes represent bootstrap values > 80% (bootstrap values were generated from 1000 iterations) and posterior probability >0.8 in the Maximum likelihood (A) and Bayesian inference (B), respectively. Both trees were rooted in bacteria and the scale bar corresponds to 0.1 substitutions per amino acid position. The values highlighted in red represent the robust support for the phylogenetic affiliation between eukaryotes and heimdallarchaeial lineages within Promethearchaeota.

## Discussion

### Expansive diversity of Promethearchaeota lineages from ancient permafrost

Our coupling of a DNA repair procedure and metagenomic sequencing resulted in successful genomic reconstruction of diverse Promethearchaeota (22 MAGs, encompassing *Heimdallarchaeia*, *Gerdarchaeia*, *Lokiarchaeia, Helarachaeia* and *Thorarchaeia*) from three depths of a single core drilled from Middle Pleistocene sediments (Kon’kovaya suite) along the East Siberian Sea coast (Figure S1). These MAGs represent novel species or genera within Promethearchaeota according to 16S rRNA similarity (Figure S3) and AAI values (Figure S5-S7). All Promethearchaeota MAGs obtained from the perennially frozen marine sediments show similarities to different archaeal lineages typically recovered from coastal or deep-sea sediments of different geographic origins (5, 9, 12, 13, 15, 16). Therefore, these ancient permafrost MAGs do not support a monophyletic radiation into the permafrost environment. Furthermore, maximum-likelihood and Bayesian analyses of the permafrost Promethearchaeota lineages provided robust support for a two-domain tree (Figure 6). The close evolutionary linkage between eukaryotes and Heimdallarchaeial lineages (Figure 6) from this expanded set of permafrost Promethearchaeota MAGs, therefore, is consistent with previous findings based on either a few (5, 8, 9, 17, 23) or much more non-permafrost Promethearchaeota lineages(6). In this regard, the prevalence of phylogenetically diverse Promethearchaeota lineages, particularly Lokiarchaeia (Figure 1), suggests that the genomes of normal marine sediment Promethearchaeota have persisted in ancient marine deposits frozen over 100 kyr ago.

### Metabolic similarity between permafrost and non-permafrost Promethearchaeota lineages

The metabolic similarity between permafrost and non-permafrost Promethearchaeota is evident in their shared heterotrophic lifestyle and fermentation capabilities (Figure 3). Consistent with previous findings from non-permafrost environments, Promethearchaeota from frozen marine sediments appear to primarily be heterotrophs that can break down carbohydrates and detrital proteins. The potential metabolic capability of the Promethearchaeota for producing formate and acetate might have partially contributed to the high concentration of these two low-molecular organic acids throughout the Kon’kovaya suite (31, 33). The identification of the Wood-Ljungdahl pathway from nearly all Lokiarchaeota MAGs (Figure 3) agreed with previous findings that fermentation of organic substrates can be coupled with the Wood–Ljungdahl carbon fixation in Lokiarchaeota (59, 60). Additionally, the presence of ferredoxin-reducing oxidoreductases (AOR, VOR, and IOR) highlights their conserved role in anaerobic biodegradation of detrital proteins in marine sediments [63]. Most importantly, pangenomic analyses of all encoded genes showed no significant gene enrichment from permafrost Promethearchaeota MAGs (Figures S9-10 and Table S5) and thereby reinforce the notion that Promethearchaeota MAGs from frozen marine sediments are metabolically similar to other Promethearchaeota from non-frozen marine sediments. Overall, all Promethearchaeota MAGs recovered from the frozen marine deposits are metabolically similar to the inferred metabolism of Promethearchaeota lineages from other environments (61). Therefore, our results highlight that certain normal marine sediment Promethearchaeota are metabolically capable of surviving under subzero temperatures over geological time.

### Evidence for survival of Promethearchaeota in ancient permafrost

Multiple lines of evidence support the survival of Promethearchaeota in frozen marine sediments over geological timescales. First, we found clear differences between the Promethearchaeota MAGs recovered from the extracellular and intracellular DNA fractions. The eDNA appeared severely damaged in the 14.8 m sediment (33), and the completeness of 21 MAGs was greatly improved by DNA repair (Figure 2 and Table S3). Therefore, MAGs from the eDNA fraction likely represent past Promethearchaeota lineages that died at some point over the past 100 kyr. This contrasts with the iDNA fraction in the 14.8 m sediment where DNA repair had minimal impact on the completeness (0-4.82%) and size (0.04-0.2 Mb) of six Promethearchaeota MAGs (Figure 2 and Table S3-S4).

Second, the relative abundances of DNA belonging to these six Promethearchaeota MAGs were much higher than other Promethearchaeota MAGs from all three depths (Figure S2). We therefore explored the possibility that the Promethearchaeota MAGs remained intact for over 100,000 years because they initially existed in greater abundance, resulting in more of their genomes surviving despite a fraction of their genomes had been degraded. The decay rate of DNA has been empirically shown to conform to an Arrhenius distribution(62), represented by the equation:

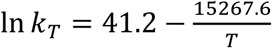

At an *in situ* permafrost temperature of -8°C [33], this results in a decay constant of k_T_=7.44×10^−8^ per base per year. Assuming a Poisson distribution of decaying genomes characterized by a whole genome decay rate (k_WG_) defined as:

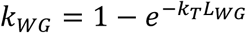

where L_WG_ represents the length of the whole genome, the fraction of genomes expected to survive over 100,000 years at -8°C can be modeled using exponential decay:

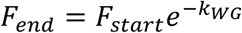

Assuming an Promethearchaeota genome size of 3 Mb, the calculated k_WG_ is approximately 0.2 genomes per year. Consequently, even if a genome initially constituted 100% of the microbial genomes, it would decline to nearly 0% after just 100 years, much less than the over 100 kyr that these sediments have been frozen. Therefore, the finding that a large abundance of MAGs from these Promethearchaeota lineages are intact with up to 92% completeness is compelling evidence that they survived because they were being maintained in living organisms.

Third, previous evidence supporting the metabolic viability of these Promethearchaeota is provided by the observation that the D/L ratio of aspartic acid from cellular proteins in these samples was maintained well lower than would be expected from abiotic racemization from dead organisms [33]. Consequently, these highly abundant Promethearchaeota represent the most promising candidates for the organisms who have partially contributed to the observed chirality of amino acids. The simplest explanation for this phenomenon is that these six Promethearchaeota lineages have survived under frozen conditions over geological time due to their adaptation to the cryogenic environment.

### Prevalence of ancient repair enzymes from Promethearchaeota and their evolutionary history

The long-term survival of microorganisms in cryogenic conditions would require active repair of damage to cellular macromolecules such as DNA and proteins. Our previous studies have implicated that the *PIMT* enzyme might be involved in maintaining low ratio of D/L aspartic acid in cellular proteins in ancient permafrost up to 3 Ma old (31, 33). Most of our Promethearchaeota MAGs harbor genes encode *PIMT* (8 MAGs) as well as methionine sulfoxide reductases (9 for *MsrA* and 7 for *MsrB*) involved in the repair of oxidatively damaged proteins (Figure S8). Furthermore, genes encoding various DNA repair enzymes (*UNG*, *MutS*, *MutL*, *RadA*, *alkB* and *UvrAB*) were also identified from our Promethearchaeota MAGs (Figure S8). In other studies, active DNA repair processes at subzero temperatures have been confirmed in microorganisms buried in ancient permafrost (33, 51). Additionally, most Promethearchaeota MAGs harbor genes potentially involved in adaptation to cold, osmotic and oxidative stresses (Figure S8). Therefore, the Promethearchaeota lineages encode various genetic machineries that would be essential for maintenance during long-term survivability under frozen conditions. Most of these genes are identified in all Promethearchaeota MAGs from permafrost sediments (Figure S8) regardless of their completeness relative to their completeness post-DNA repair. Moreover, these genes are also present in various Promethearchaeota lineages from non-permafrost environments. This suggests that the survival mechanisms for long-term survivability under cryogenic conditions are a common feature across Promethearchaeota lineages (Figure 2).

DNA repair mechanisms of the Promethearchaeota have previously been shown to be shared with the later-arising eukaryotes, following their phylogenetic relationships(63). In contrast, we found that the PIMT protein from the Promethearchaeota lineages is closely related to the TACK group instead of eukaryotes, differing from their vertical inheritance patterns. Likewise, the evolutionary histories of *MsrA* and *MsrB* (Figure 5) support our finding that protein repair strategies in Promethearchaeota were acquired through multiple lateral gene transfer events, rather than the vertical eukaryotic lineage. The genes *MsrA* and *MsrB* encode two structurally distinct enzymes for repairing the oxidative damage of methionine residues caused by reactive oxygen species (30). Due to unequal rates of substitution and frequent horizontal gene transfer events (64), the complex evolutionary history *MsrA* and *MsrB* from the Promethearchaeota suggested that Promethearchaeota MAGs likely acquired *MsrA* and *MsrB* from bacteria through multiple ancient horizontal gene transfers. Protein repair dominates microbial energy expenditures under severe maintenance conditions(28). This may explain why Promethearchaeota protein repair mechanisms appear to be under different selective pressure than their purely vertically inherited DNA repair mechanisms.

## Conclusions

The integration of laboratory-based DNA repair techniques and metagenomic sequencing allowed us to successfully reconstruct 15 medium- to high-quality Promethearchaeota MAGs from marine sediments frozen 100-120 kyr ago. Many were of high completeness even without applying protein repair mechanisms, suggesting that they survived being frozen for this long. However, they had similar phylogenies and metabolisms (fermentation of carbohydrates and detrial proteins) to a wide range of Promethearchaeota found in non-frozen subsurface environments. Phylogenetic analyses of ancient protein repair enzymes indicated that the genes involved in the repair of isomerized/racemized aspartic acid (*PIMT*) and oxidized methionine residues (*MsrA* and *MsrB*) were horizontally transferred independently of the divergence of eukaryotes. The resilience and survival potential of ancient Promethearchaeota cells in long-term cryogenic conditions hints at the possibility of similar adaptations of these eukaryogenesis-related archaea in extreme cold environments on other planets such as Mars and Europa. Our results suggest that surviving being frozen for over 100 kyr is a widespread trait among Promethearchaeota, suggesting that the phylum is well-adapted to survive over geological timescales while maintaining a slow metabolic activity sufficient to repair proteins and DNA.

## Acknowledgments

This research is dedicated to the memory of TC Onstott, without whom this work would not be possible, including securing funding, providing guidance, and working on earlier versions of this manuscript. This research was supported by an NSF DEB-1442059 and NSF EAR-1528492 to TCO, NSF DEB-1442262 to TAV, TCO and KGL, and NSF International Research Experience for Students grant IIA-1460058 to TAV and KGL, U.S. Department of Energy, Office of Science, Office of Biological and Environmental Research, Genomic Science Program under Award Number DE-SC0020369 to KGL, TAV, and TCO and by Russian Government Assignment AAAA-A18-118013190181-6 and RFBR 19-29-05003 to EMR. Authors thank undergraduate student Molly Moran, participant of the NSF International Research Experience for Students project, for technical help.

