## Supplementary material for "Traits enabling persistence of living Promethearchaeota in marine sediments frozen for over 100 kyr": Tables S1-4 and Figures S1-10

Mailing address: No. 68 Jincheng Street, East Lake High-tech Development Zone, Wuhan, Hubei Province, China, 430078

Table S1 Statistical summary for all Asgard MAGs recovered from the Kon'kovaya suite marine horizon.

| MAGs | Completeness (%) | Contamination (%) | GC (%) | L50 | N50  (kb) | Size (Mb) | Number of scaffolds |
| --- | --- | --- | --- | --- | --- | --- | --- |
| Heimdallarchaeia_Ch1_17_bin51 | 93.22 | 3.27 | 33.8 | 75 | 12.1 | 3.09 | 366 |
| Gerdarchaeia_Ch1_17_bin89 | 91.59 | 7.94 | 34.1 | 16 | 63.4 | 3.36 | 115 |
| Lokiarchaeia_Ch1_17_bin106 | 88.79 | 2.34 | 26.8 | 45 | 21.2 | 3.36 | 288 |
| Thorarchaeia_Ch1_17_bin66 | 88.32 | 4.74 | 44.4 | 41 | 20.9 | 3.04 | 442 |
| Lokiarchaeia_Ch1_17_bin60 | 88.32 | 5.14 | 31.6 | 20 | 62.8 | 3.92 | 175 |
| Lokiarchaeia_Ch1_17_bin48 | 84.51 | 3.27 | 32.2 | 50 | 14.8 | 3.19 | 403 |
| Lokiarchaeia_Ch1_17_bin8 | 83.33 | 4.21 | 31.6 | 34 | 28.1 | 3.52 | 225 |
| Lokiarchaeia_Ch1_17_bin12 | 81.31 | 2.34 | 31.4 | 51 | 16.7 | 2.77 | 244 |
| Lokiarchaeia_Ch1_17_bin49 | 78.9 | 2.8 | 32.3 | 30 | 26.6 | 2.53 | 170 |
| Lokiarchaeia_Ch1_17_bin22 | 75.78 | 3.02 | 31.5 | 69 | 9.4 | 2.14 | 295 |
| Lokiarchaeia_Ch1_17_bin41 | 73.14 | 3.74 | 31.9 | 59 | 15.8 | 3.22 | 279 |
| Lokiarchaeia_Ch1_17_bin40 | 69.09 | 5.37 | 31.6 | 59 | 11.4 | 2.39 | 585 |
| Lokiarchaeia_Ch1_17_bin33 | 65.42 | 2.4 | 32.2 | 172 | 4.9 | 2.17 | 522 |
| Lokiarchaeia_Ch1_17_bin24 | 64.64 | 2.34 | 26.2 | 182 | 3.6 | 2.04 | 625 |
| Lokiarchaeia_Ch1_17_bin21 | 58.21 | 1.4 | 32.1 | 126 | 4.7 | 2.43 | 419 |
| Lokiarchaeia_Ch1_17_bin44 | 40.31 | 7.48 | 30.3 | 638 | 2.1 | 3.48 | 1626 |
| Thorarchaeia_Ch1_17_bin23 | 39.34 | 4.21 | 43.1 | 73 | 6.9 | 1.88 | 418 |
| Lokiarchaeia_Ch1_17_bin9 | 38.7 | 5.14 | 32.2 | 213 | 1.9 | 1.11 | 598 |
| Thorarchaeia_Ch1_17_bin34 | 37.54 | 4.91 | 42.9 | 48 | 8.2 | 1.29 | 197 |
| Lokiarchaeia_Ch1_17_bin7 | 37.02 | 3.27 | 30.1 | 36 | 7.9 | 0.83 | 155 |
| Lokiarchaeia_Ch1_17_bin37 | 33.8 | 1.4 | 31.4 | 57 | 8.2 | 1.33 | 195 |
| Helarchaeia_Ch1_17_bin36 | 14.09 | 0 | 38.7 | 54 | 2.4 | 0.38 | 155 |

**Table S2** Taxonomic classification of all Asgard MAGs based on the GTDB-Tk tool.

| MAGs | Taxonomic classification |
| --- | --- |
| Heimdallarchaeia_Ch1_17_bin51 | d__Archaea;p__Asgardarchaeia;c__Heimdallarchaeia;o__UBA460;f__UBA460;g__;s__ |
| ***Gerdarchaeia***_Ch1_17_bin89 | d__Archaea;p__Asgardarchaeia;c__Heimdallarchaeia;o__;f__;g__;s__ |
| Thorarchaeia_Ch1_17_bin23 | d__Archaea;p__Asgardarchaeia;c__Thorarchaeia;o__Thorarchaeales;f__Thorarchaeaceae;g__MP8T-1;s__ |
| Thorarchaeia_Ch1_17_bin34 | d__Archaea;p__Asgardarchaeia;c__Thorarchaeia;o__Thorarchaeales;f__Thorarchaeaceae;g__MP8T-1;s__ |
| Thorarchaeia_Ch1_17_bin66 | d__Archaea;p__Asgardarchaeia;c__Thorarchaeia;o__Thorarchaeales;f__Thorarchaeaceae;g__SMTZ1-45;s__ |
| Lokiarchaeia_Ch1_17_bin49 | d__Archaea;p__Asgardarchaeia;c__Lokiarchaeia;o__CR-4;f__SOKP01;g__SOKP01;s__ |
| Lokiarchaeia_Ch1_17_bin9 | d__Archaea;p__Asgardarchaeia;c__Lokiarchaeia;o__CR-4;f__SOKP01;g__SOKP01;s__ |
| Lokiarchaeia_Ch1_17_bin41 | d__Archaea;p__Asgardarchaeia;c__Lokiarchaeia;o__CR-4;f__SOKP01;g__SOKP01;s__ |
| Lokiarchaeia_Ch1_17_bin21 | d__Archaea;p__Asgardarchaeia;c__Lokiarchaeia;o__CR-4;f__SOKP01;g__SOKP01;s__ |
| Lokiarchaeia_Ch1_17_bin40 | d__Archaea;p__Asgardarchaeia;c__Lokiarchaeia;o__CR-4;f__SOKP01;g__SOKP01;s__ |
| Lokiarchaeia_Ch1_17_bin48 | d__Archaea;p__Asgardarchaeia;c__Lokiarchaeia;o__CR-4;f__SOKP01;g__SOKP01;s__ |
| Lokiarchaeia_Ch1_17_bin33 | d__Archaea;p__Asgardarchaeia;c__Lokiarchaeia;o__CR-4;f__SOKP01;g__SOKP01;s__ |
| Lokiarchaeia_Ch1_17_bin12 | d__Archaea;p__Asgardarchaeia;c__Lokiarchaeia;o__CR-4;f__SOKP01;g__SOKP01;s__ |
| Lokiarchaeia_Ch1_17_bin22 | d__Archaea;p__Asgardarchaeia;c__Lokiarchaeia;o__CR-4;f__SOKP01;g__SOKP01;s__ |
| Lokiarchaeia_Ch1_17_bin7 | d__Archaea;p__Asgardarchaeia;c__Lokiarchaeia;o__CR-4;f__SOKP01;g__SOKP01;s__ |
| Lokiarchaeia_Ch1_17_bin60 | d__Archaea;p__Asgardarchaeia;c__Lokiarchaeia;o__CR-4;f__SOKP01;g__SOKP01;s__ |
| Lokiarchaeia_Ch1_17_bin8 | d__Archaea;p__Asgardarchaeia;c__Lokiarchaeia;o__CR-4;f__SOKP01;g__SOKP01;s__ |
| Lokiarchaeia_Ch1_17_bin37 | d__Archaea;p__Asgardarchaeia;c__Lokiarchaeia;o__CR-4;f__SOKP01;g__SDNM01;s__ |
| Lokiarchaeia_Ch1_17_bin44 | d__Archaea;p__Asgardarchaeia;c__Lokiarchaeia;o__CR-4;f__SOKP01;g__SDNM01;s__ |
| Lokiarchaeia_Ch1_17_bin106 | d__Archaea;p__Asgardarchaeia;c__Lokiarchaeia;o__CR-4;f__SOKP01;g__;s__ |
| Lokiarchaeia_Ch1_17_bin24 | d__Archaea;p__Asgardarchaeia;c__Lokiarchaeia;o__CR-4;f__SOKP01;g__;s__ |
| Helarchaeia_Ch1_17_bin36 | d__Archaea;p__Asgardarchaeia;c__Lokiarchaeia;o__Helarchaeales;f__HEL-GB-A;g__HEL-GB-A;s__ |

**Table S3** Completeness (%) of MAGs recovered from extracellular and intracellular DNA fraction with and without DNA repair for 14.8 m sample. The MAGs with minimum improvement after DNA repair were highlighted in bold.

| MAGs | 14.8e | 14.8_preCR | 14.8i | 14.8i_preCR |
| --- | --- | --- | --- | --- |
| Heimdallarchaeia_Ch1_17_bin51 | 0 | 61.91 | 33.68 | 70.95 |
| ***Gerdarchaeia*_Ch1_17_bin89** | 26.47 | 92.05 | 92.52 | 92.52 |
| Lokiarchaeia_Ch1_17_bin106 | 0 | 19.62 | 36.71 | 77.57 |
| Thorarchaeia_Ch1_17_bin66 | 0 | 37.09 | 35.31 | 73.68 |
| Lokiarchaeia_Ch1_17_bin60 | 0 | 73.7 | 64.48 | 84.57 |
| **Lokiarchaeia_Ch1_17_bin48** | 0 | 81.7 | 81.77 | 84.11 |
| Lokiarchaeia_Ch1_17_bin8 | 4.09 | 81.77 | 73.05 | 83.17 |
| **Lokiarchaeia_Ch1_17_bin12** | 0 | 78.73 | 79.67 | 81.3 |
| **Lokiarchaeia_Ch1_17_bin49** | 0 | 77.49 | 77.03 | 79.83 |
| **Lokiarchaeia_Ch1_17_bin22** | 0 | 73.44 | 70.96 | 75.78 |
| **Lokiarchaeia_Ch1_17_bin41** | 4.16 | 68.32 | 67.54 | 72.13 |
| Lokiarchaeia_Ch1_17_bin40 | 0 | 54.82 | 47.35 | 66.82 |
| Lokiarchaeia_Ch1_17_bin33 | 0 | 2.8 | 4.72 | 26.82 |
| Lokiarchaeia_Ch1_17_bin24 | 0 | 10.43 | 25.38 | 56.06 |
| Lokiarchaeia_Ch1_17_bin21 | 0 | 31.34 | 32.3 | 54.08 |
| Lokiarchaeia_Ch1_17_bin44 | 0 | 4.97 | 7.45 | 23.42 |
| Thorarchaeia_Ch1_17_bin23 | 0 | 18.22 | 13.58 | 23.75 |
| Lokiarchaeia_Ch1_17_bin9 | 0 | 15.42 | 20.08 | 35.57 |
| Thorarchaeia_Ch1_17_bin34 | 0 | 4.16 | 5.45 | 19.15 |
| Lokiarchaeia_Ch1_17_bin7 | 0 | 10.85 | 4.16 | 5.45 |
| Lokiarchaeia_Ch1_17_bin37 | 0 | 7.94 | 16.85 | 28.19 |
| Helarchaeia_Ch1_17_bin36 | 0 | 0 | 0 | 0 |

**Table S4** Genome size (Mb) of MAGs recovered from extracellular and intracellular DNA fraction with and without DNA repair for 14.8 m sample.

| MAGs | 14.8e | 14.8_preCR | 14.8i | 14.8i_preCR |
| --- | --- | --- | --- | --- |
| Heimdallarchaeia_Ch1_17_bin51 | 0.006 | 1.634 | 0.805 | 2.015 |
| Gerdarchaeia_Ch1_17_bin89 | 0.895 | 3.394 | 3.377 | 3.413 |
| Lokiarchaeia_Ch1_17_bin106 | 0.002 | 0.838 | 1.551 | 2.911 |
| Thorarchaeia_Ch1_17_bin66 | 0.002 | 0.894 | 1.058 | 2.122 |
| Lokiarchaeia_Ch1_17_bin60 | 0.019 | 2.808 | 2.878 | 3.794 |
| Lokiarchaeia_Ch1_17_bin48 | 0.076 | 2.992 | 3.053 | 3.284 |
| Lokiarchaeia_Ch1_17_bin8 | 0.096 | 3.197 | 2.518 | 3.32 |
| Lokiarchaeia_Ch1_17_bin12 | 0.031 | 2.567 | 2.725 | 2.825 |
| Lokiarchaeia_Ch1_17_bin49 | 0.023 | 2.391 | 2.427 | 2.536 |
| Lokiarchaeia_Ch1_17_bin22 | 0.038 | 1.953 | 1.959 | 2.163 |
| Lokiarchaeia_Ch1_17_bin41 | 0.254 | 3.109 | 3.019 | 3.226 |
| Lokiarchaeia_Ch1_17_bin40 | 0.009 | 1.465 | 1.521 | 2.19 |
| Lokiarchaeia_Ch1_17_bin33 | 0 | 0.113 | 0.082 | 0.774 |
| Lokiarchaeia_Ch1_17_bin24 | 0.001 | 0.288 | 0.851 | 1.954 |
| Lokiarchaeia_Ch1_17_bin21 | 0.001 | 0.894 | 1.146 | 2.138 |
| Lokiarchaeia_Ch1_17_bin44 | 0.003 | 0.306 | 0.28 | 1.569 |
| Thorarchaeia_Ch1_17_bin23 | 0.007 | 0.487 | 0.299 | 0.719 |
| Lokiarchaeia_Ch1_17_bin9 | 0.002 | 0.41 | 0.409 | 1.076 |
| Thorarchaeia_Ch1_17_bin34 | 0 | 0.289 | 0.223 | 0.596 |
| Lokiarchaeia_Ch1_17_bin7 | 0 | 0.075 | 0.067 | 0.252 |
| Lokiarchaeia_Ch1_17_bin37 | 0 | 0.161 | 0.476 | 1.113 |
| Helarchaeia_Ch1_17_bin36 | 0 | 0.032 | 0.007 | 0.046 |

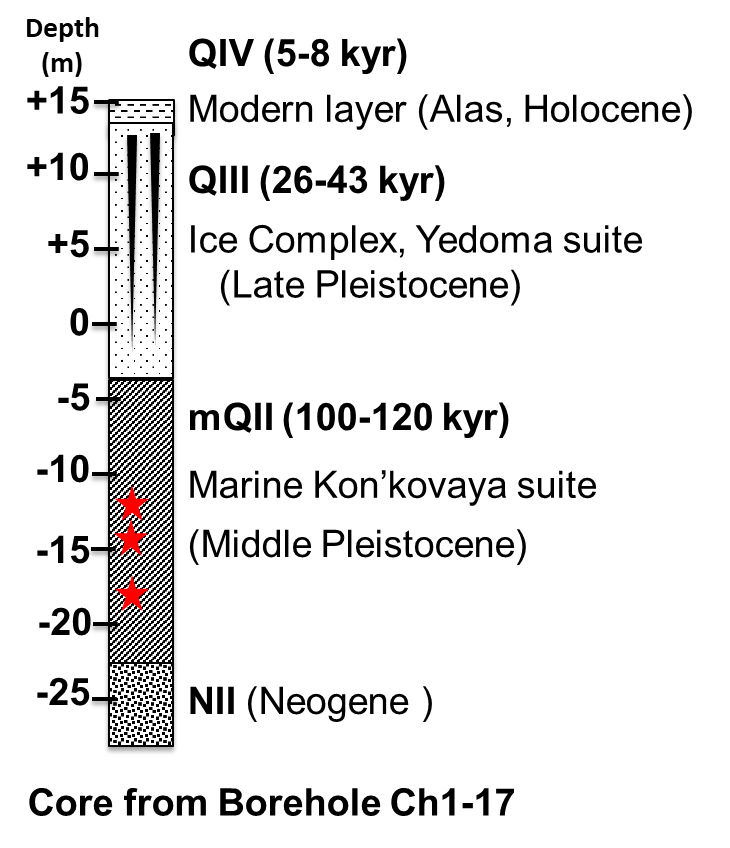

**Figure S1** A schematic of the drilled the core sediment (~22 m) from borehole Ch1-17 at Cape Chukochii near the East Siberian Sea coast. The red stars indicate the depth of sediments (13.4, 14.8 and 18.3 m, meters below land surface) that were selected for metagenomic sequencing. Scales above 0 refer to the adjacent crops above the borehole.

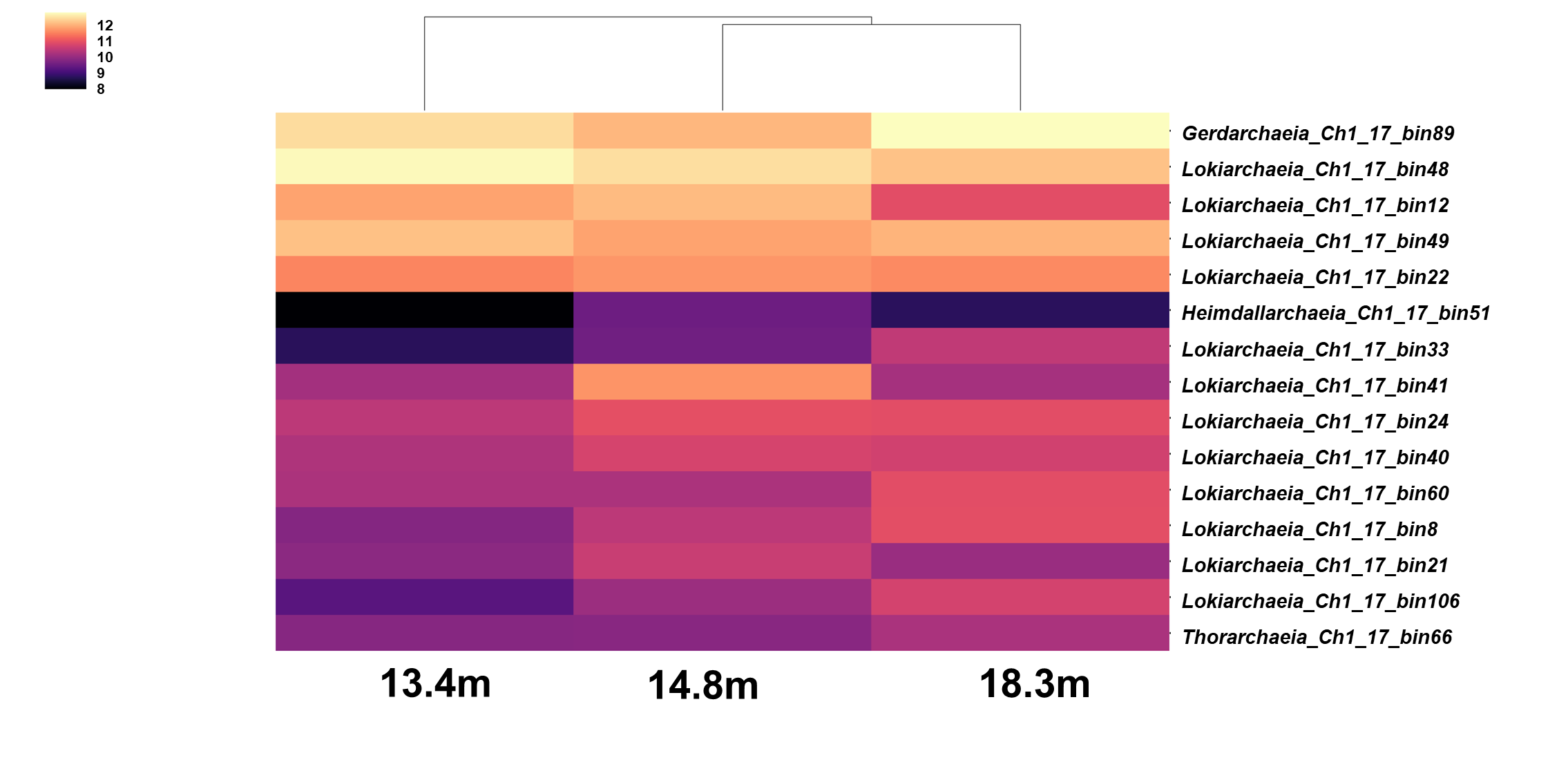

**Figure S2** Relative abundance of MAGs across the three metagenomes (13.4, 14.8 and 18.3m). The abundance of each MAG was calculated from the reads mapped in the metagenome and normalized to the individual sample size as genome copies per million reads.

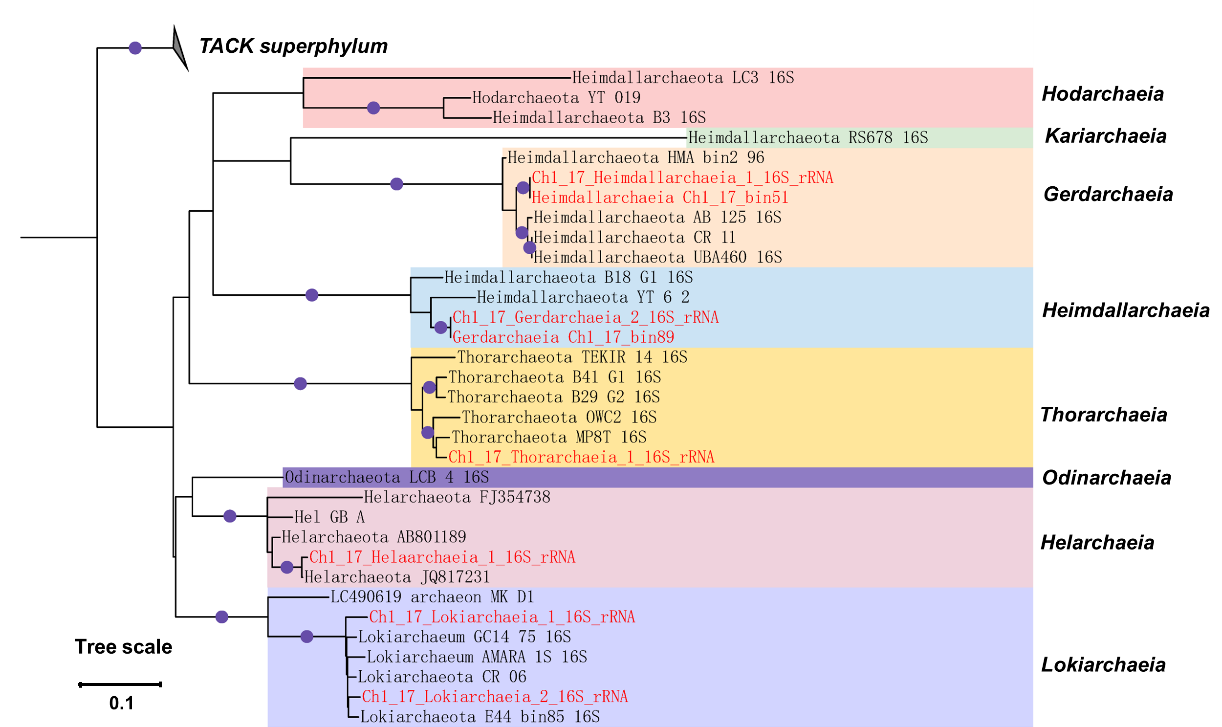

**Figure S3** Phylogenetic tree based on 16S rRNA genes from Asgard MAGs or metagenomes and close relatives of culture and uncultured organisms. The purple dots represent bootstrap values > 70% (bootstrap values were generated from 1000 replications). The 16S rRNA genes from MAGs in this study were highlighted in bold and red. The TACK group was used as an outgroup. The scale bar corresponds to 0.1 substitutions per nucleic acid position.

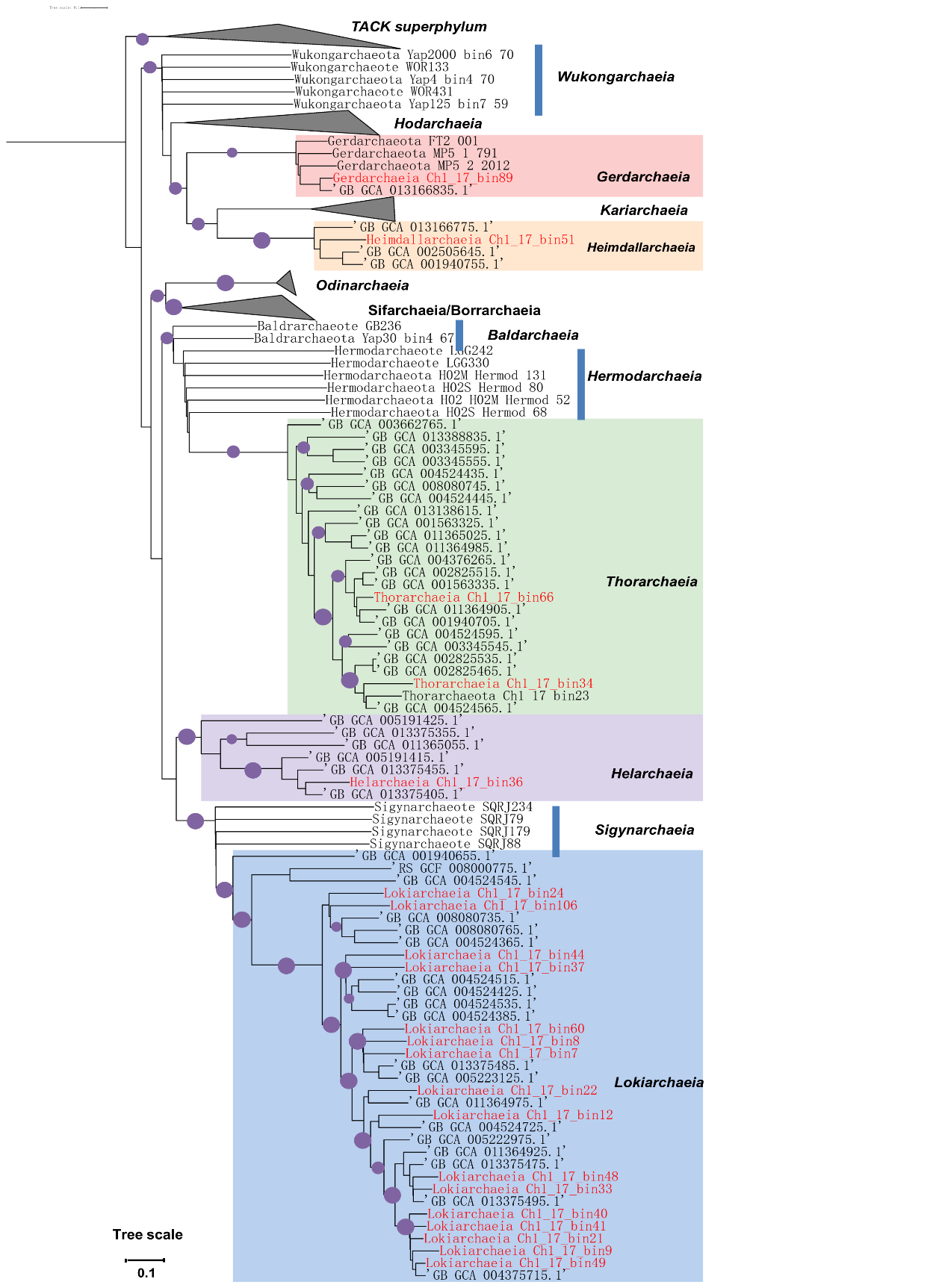

**Figure S4** Maximum-likelihood phylogenomic tree of all reconstructed Asgard MAGs inferred from concatenated 122 single-copy marker genes using the GTDB-Tk toolkit. The purple dots represent bootstrap values > 70% (bootstrap values were generated from 1000 replications). The MAGs in this study were highlighted in bold and red and the TACK group was used as an outgroup. The scale bar corresponds to 0.1 substitutions per amino acid position.

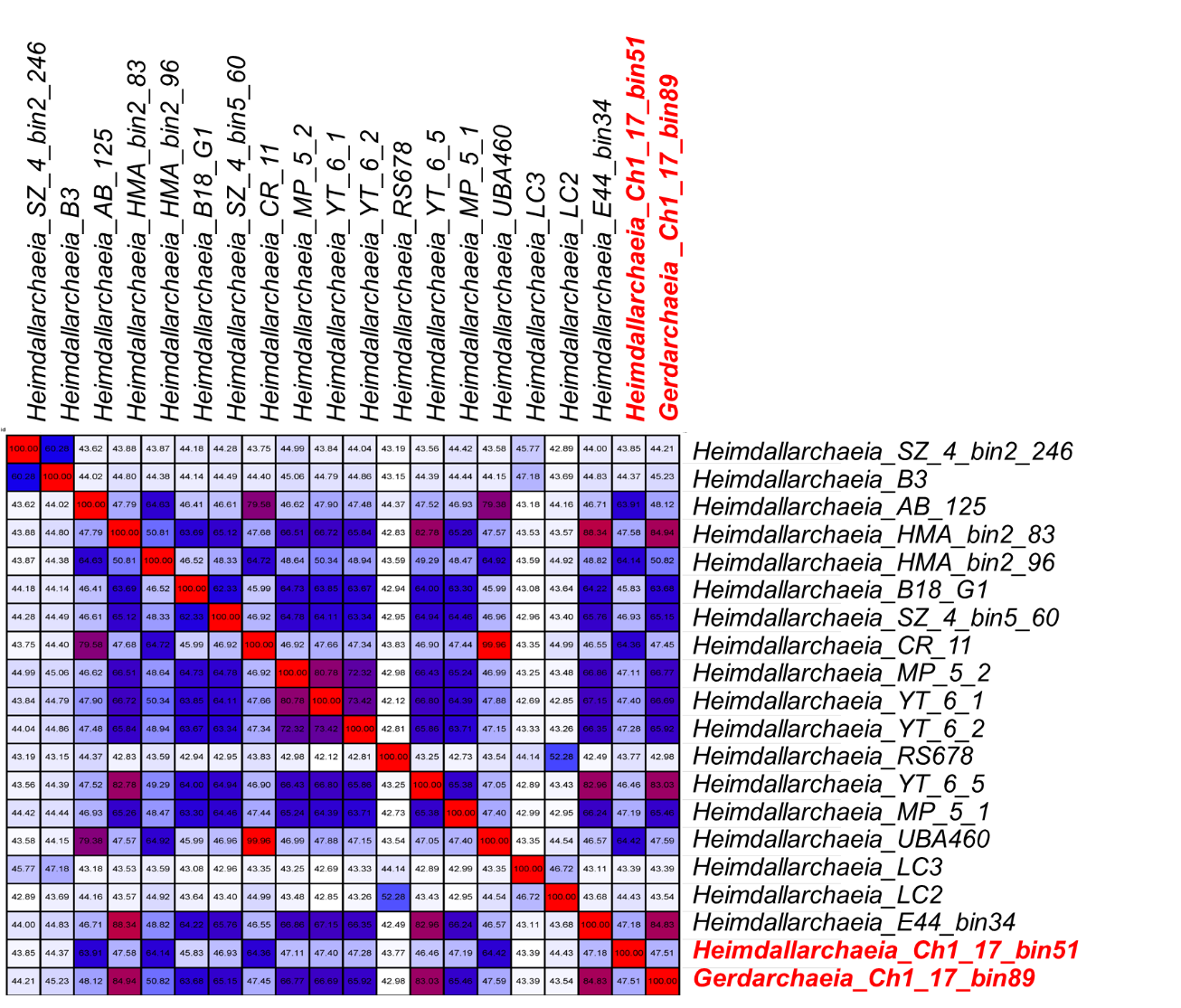

**Figure S5** Pairwise average amino acid identity (AAI) distances among the *Heimdallarchaeia*-related MAGs and their closest genomic relatives. The MAGs related to *Heimdallarchaeia* and *Gerdarchaeia* from this study were highlighted in bold and red.

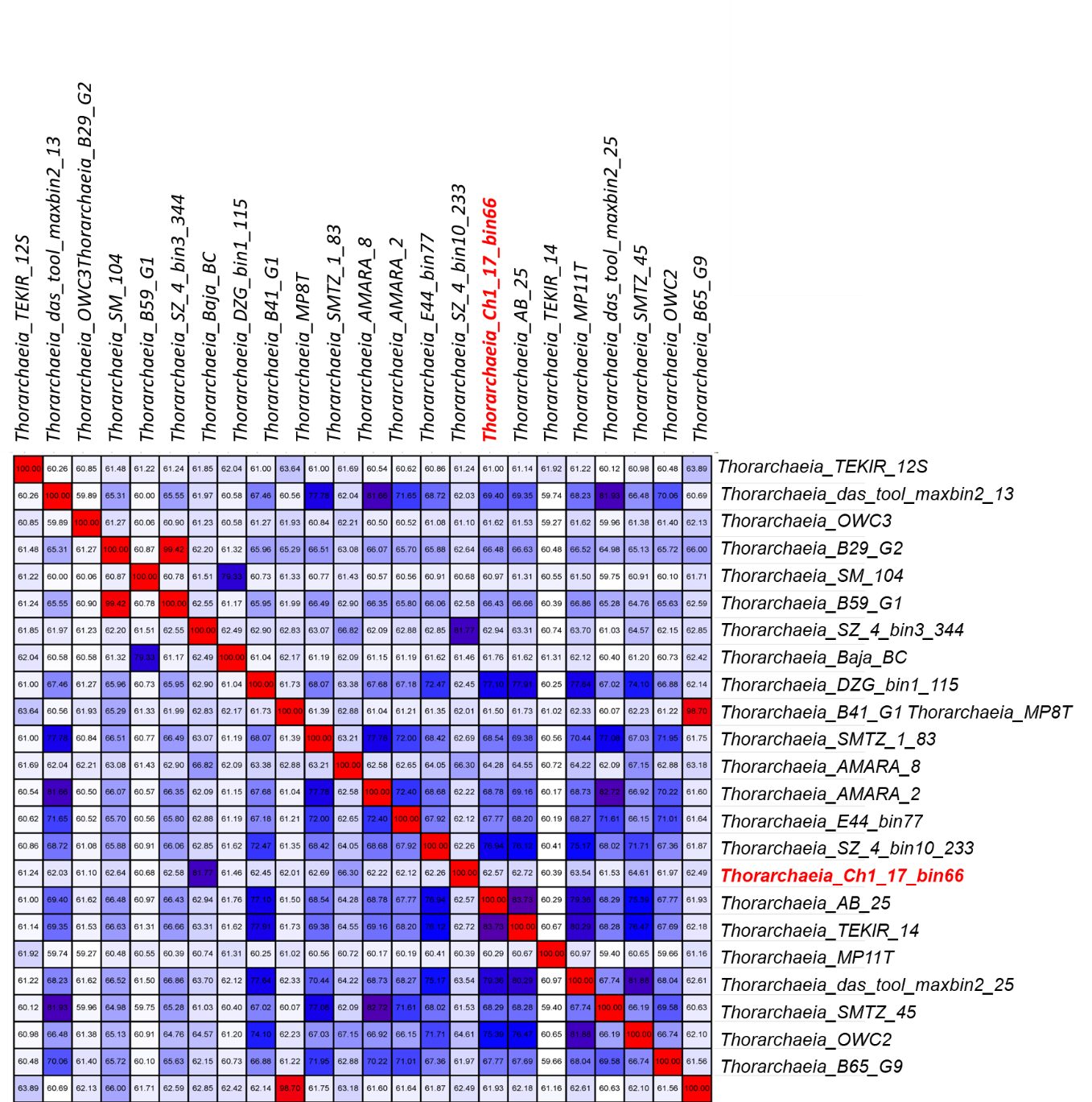

**Figure S6** Pairwise average amino acid identity (AAI) distances among the *Thorarchaeia*-related MAGs and their closest genomic relatives. The *Thorarchaeia*-related MAGs in this study were highlighted in bold and red

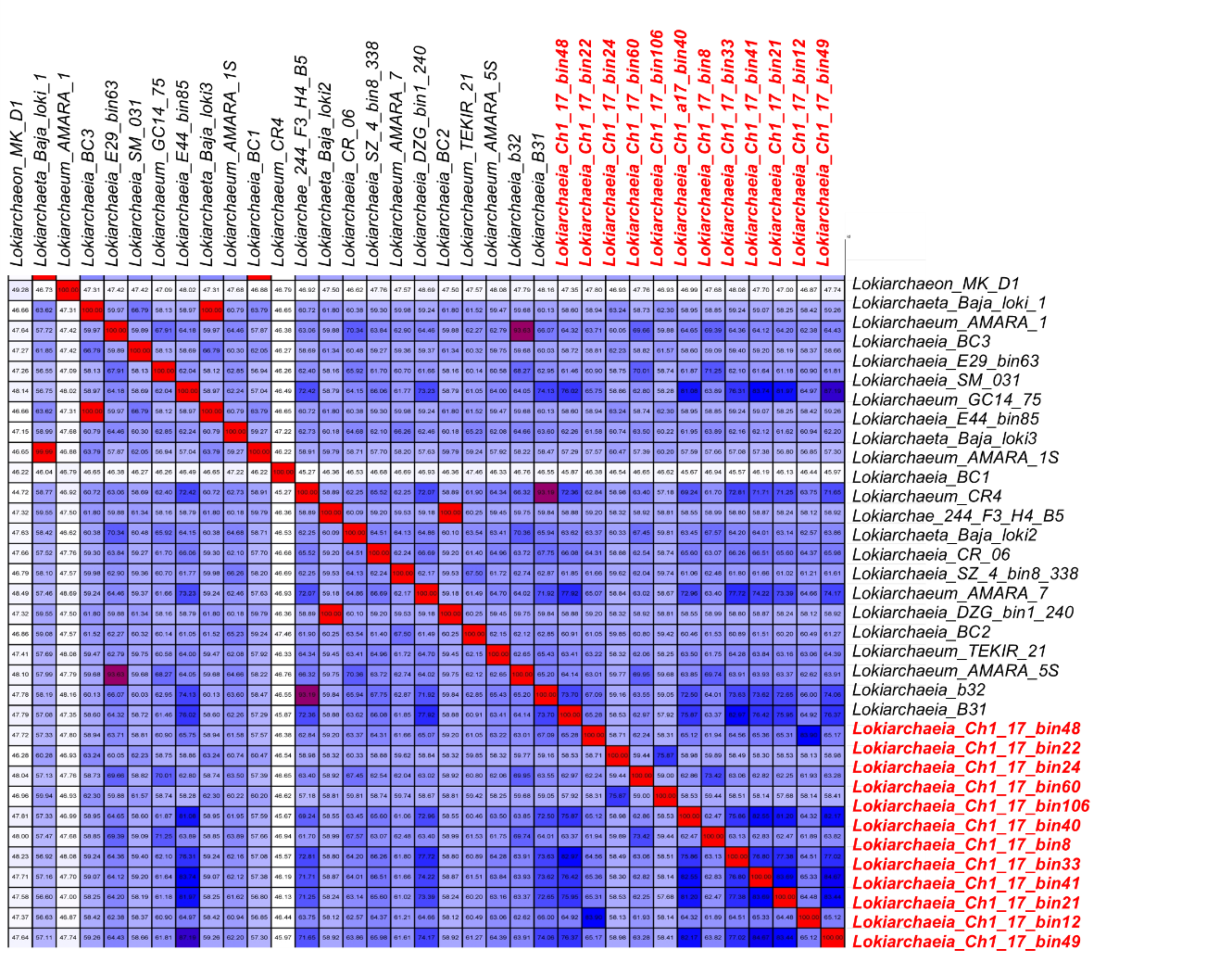

**Figure S7** Pairwise average amino acid identity (AAI) distances among the *Lokiarchaeia*-related MAGs and their closest genomic relatives. The *Lokiarchaeia*-related MAGs in this study were highlighted in bold and red.

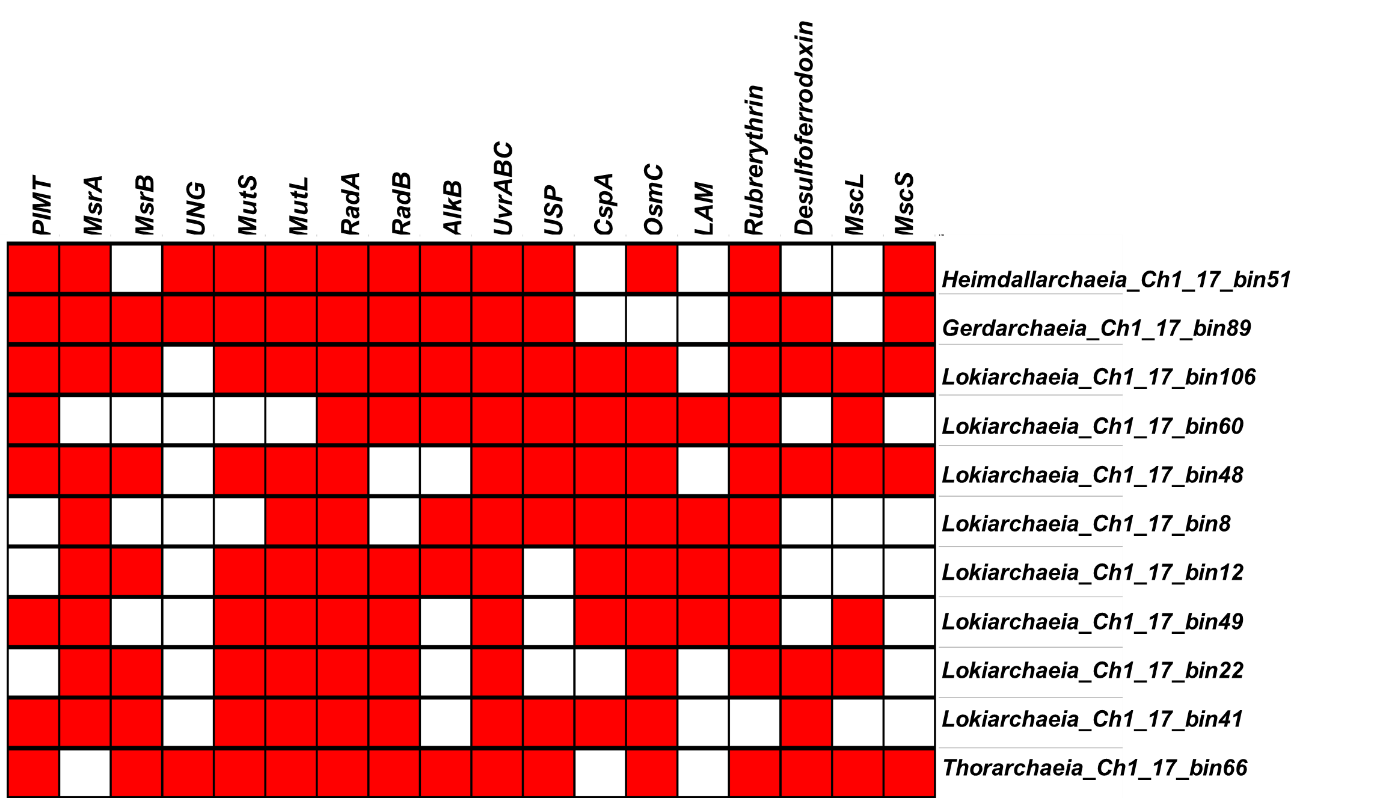

**Figure S8** Presence (red) of functional genes in Asgard MAGs that might be potentially involved in the repair of DNA and protein damage and survival under cold, osmotic and oxidative stresses. Gene abbreviations: protein L-isoaspartyl/D-aspartyl o-methyltransferase (*PIMT*), methionine sulfoxide reductase (*MsrA*), uracil-DNA glycosylase (*UNG*), DNA mismatch repair protein (*MutS*), DNA repair proteins (*RadA and RadB*), DNA alkylation repair protein *(alkB*), *UvrABC* endonuclease (*UvrABC*), Universal stress protein (*USP*), cold shock protein (*CspA*), Osmotically inducible protein (*OsmC*), lysine 2,3-aminomutase (*LAM*), Large-conductance mechanosensitive channel (*MscL*) and Small-conductance mechanosensitive channel (*MscS*).

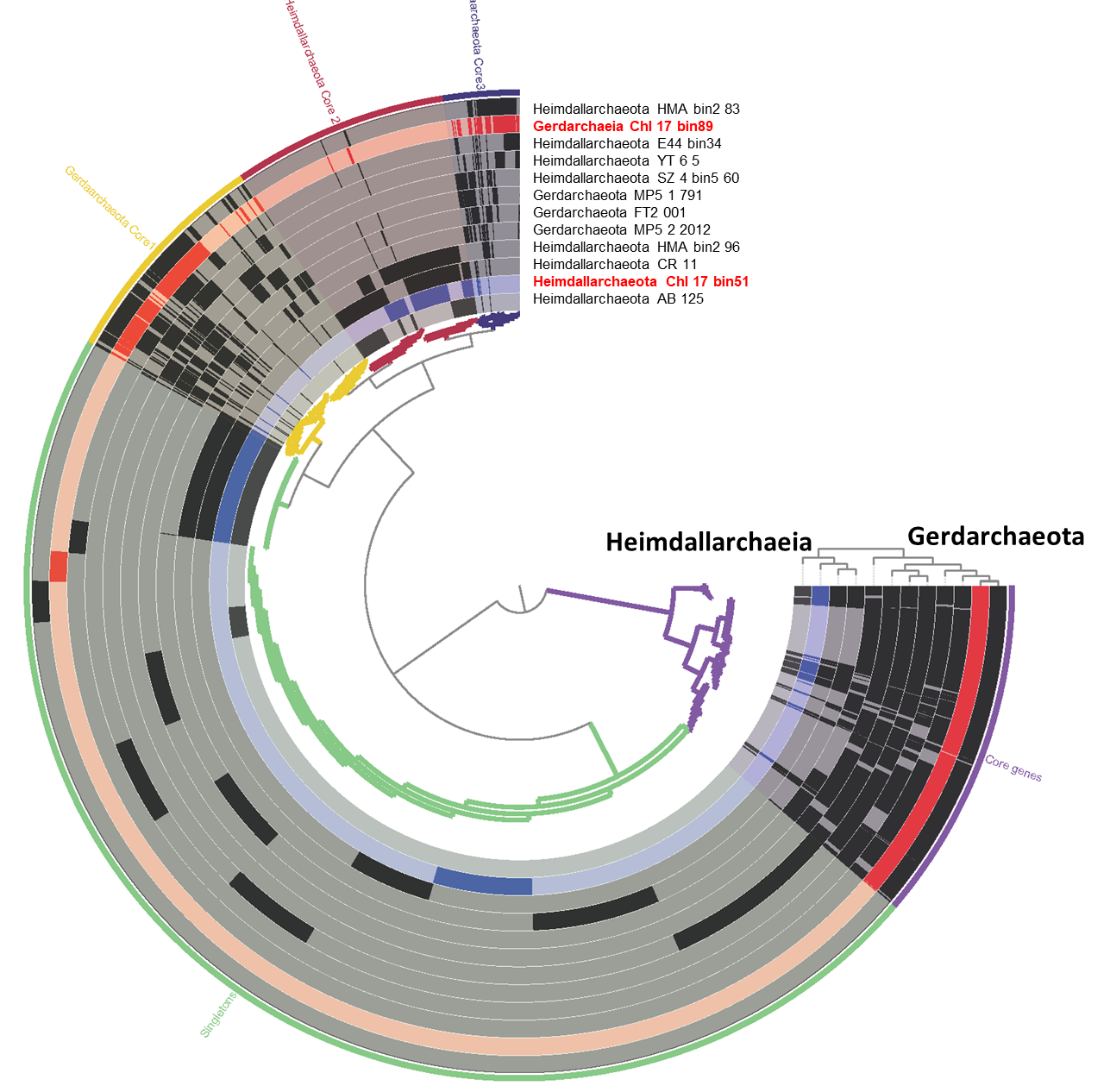

**Figure S9** Pangenomics of Heimdallarchaeia and Gerdarchaeia MAGs from ancient permafrost (highlighted in red) and other environments. The core gens and singletons are shown in the outmost layer.

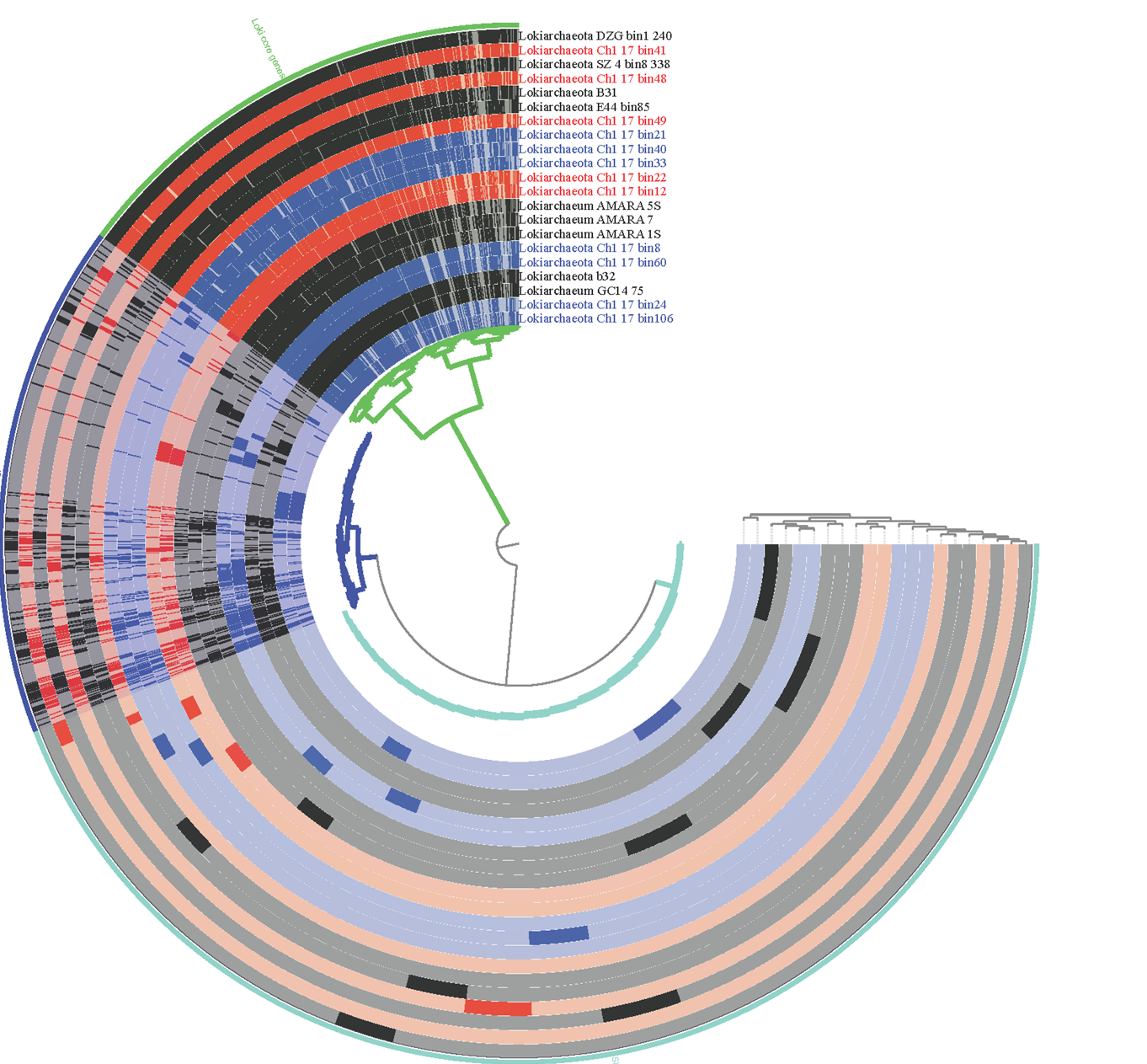

**Figure S10** Pangenomics of Lokiarchaeia MAGs from ancient permafrost (highlighted in red) and other environments. The core gens and singletons are shown in the outmost layer.
